# Ciliary ARL13B drives renal cystogenesis via its GEF activity

**DOI:** 10.1101/2025.11.20.689124

**Authors:** Robert E. Van Sciver, Avery Forster, Leslie M. Lewis, Tamara Caspary

## Abstract

**Background:** Polycystic kidney disease (PKD) is the leading genetic cause of renal failure, resulting in the accumulation of fluid filled cysts and gross enlargement of the kidney. Mutations in *PKD1* or *PKD2*, which encode ciliary polycystin proteins, are the most common cause of PKD. These proteins function in a cilia-dependent cyst activation (CDCA) pathway–one that requires cilia for its pro-cystic function–yet the molecular driver(s) of this pathway are unknown. ARL13B is a regulatory GTPase enriched in cilia and links to renal cystogenesis. ARL13B possesses guanine nucleotide exchange factor (GEF) activity for ARL3, another ARL with links to cilia.

**Methods:** We used two distinct *Arl13b* mouse alleles to investigate whether ARL13B is a component of the CDCA pathway: *Arl13b^V358A^* encodes for enzymatically normal ARL13B that is undetectable in cilia, and *Arl13b^R79Q^* encodes for cilia-localized ARL13B lacking a residue critical for its ARL3 GEF activity. We used these alleles in a *Pkd1*-deficient adult mouse model and investigated renal morphology (H&E and cystic index analysis), physiology (blood urea nitrogen measurements), renal fibrosis (picrosirius staining and α-smooth muscle actin levels), renal injury (SOX9 immunofluorescent staining and quantification), and Wnt signaling (β-catenin and cyclin D1 protein levels).

**Results:** We found that loss of ciliary ARL13B or mutation of a single residue critical for its ARL3 GEF activity suppressed *Pkd1*-dependent cysts. We observed reductions in kidney size, cystic index, and blood urea nitrogen. We also observed suppression of renal fibrosis, renal injury, and β-catenin and cyclin D1 protein levels.

**Conclusions:** Our results identified a subcellular location and mechanism driving *Pkd1*-dependent renal cystogenesis. We demonstrated that expression of a critical residue for ARL13B’s GEF activity specifically in cilia is a key mechanism of the CDCA pathway driving renal cystogenesis.

**Key Points:**

- Loss of ciliary ARL13B suppressed renal cystogenesis in an adult mouse model of polycystic kidney disease (PKD) without ablating cilia
- Loss of ciliary ARL13B or mutation of the residue critical for its GEF activity did not affect renal morphology or physiology in an adult induced mouse model
- Mutation of a residue critical for ARL13B activation of ARL3 suppressed cystic phenotypes in *Pkd1*-dependent cysts

## Introduction

Autosomal dominant polycystic kidney disease (ADPKD) is the most common monogenic disease and the leading genetic cause of renal failure, affecting an estimated 1:500 to 1:1000 people worldwide^1–4^. Mutations in either *PKD1* or *PKD2* account for over 95% of cases of ADPKD^5–7^. These genes encode the polycystin 1 (PC1) and polycystin 2 (PC2) proteins, which localize to the primary cilium^8–10^. Primary cilia are microtubule-based organelles found on nearly every cell; they provide a privileged compartment linked to several signaling pathways^11–15^. In the kidney, cilia protrude into the lumen of the nephron where they may sense and respond to lumen fluid flow^16–19^.

Loss of polycystin function or exclusion of polycystin proteins from cilia causes severe renal cyst formation^20,21^. A multitude of aberrant signaling pathways and pathologies including Wnt/β-catenin/cyclin D1 signaling, renal injury, and fibrosis accompany cyst formation in PKD patient samples and animal models^22–33^. Genetic mouse models show that targeting these pathways relieves cyst progression and restores normal signaling^28,34–41^. Loss of centrosomal proteins, found at the base of cilia, or loss of cilia themselves leads to suppression of severe cystic phenotypes^37,40–42^. The most parsimonious interpretation of these results is the hypothesis of a cilia-dependent cyst activating (CDCA) pathway^37,43,44^. Simultaneous loss of cilia and polycystins suppresses the severe cysts caused by loss of polycystins alone, arguing that polycystin proteins normally function within cilia to inhibit this yet-to-be-identified pro-cystic pathway^37,43,44^. Components of the CDCA pathway are predicted to localize to cilia, and their intra-cilial loss is expected to suppress polycystin-dependent renal cysts. While discovery-based approaches have identified regulators and targets of the CDCA pathway, the question of the molecular identity of any ciliary drivers of the CDCA pathway remains unanswered^34^. Identifying the molecular components of this pathway will likely identify novel molecular targets for therapy in ADPKD^45^.

ARL13B is an ADP-ribosylation factor-like (ARL) family regulatory GTPase enriched in primary cilia whose loss in the kidney causes cysts^46–49^. Loss of *Arl13b* in kidney also leads to a loss of renal cilia^48,49^. In addition to its GTPase activity, ARL13B serves as a guanine nucleotide exchange factor (GEF) for ARL3, another ARL family GTPase that can localize to primary cilia^50,51^. Our work on ARL13B identified mutations that allow us to separate its cilia localization from its known GEF activity^51–57^. *Arl13b^V358A^* is an engineered gene variant that disrupts the ARL13B RVxP cilia localization motif^52–55^. ARL13B^V358A^ protein is stably expressed and retains GTP binding and GEF activities but is undetectable in cilia^53–55^. *Arl13b^V358A/V358A^* mice display normal ciliation and cilia length in their kidneys, however they develop mild cystogenesis during development, suggesting ARL13B exhibits distinct ciliary and non-ciliary functions in the kidney^55^. In contrast, the ARL13B^R79Q^ mutation abolishes ARL13B GEF activity for ARL3 *in vitro* without altering its cilia localization, and mice homozygous for this mutation do not develop kidney cysts^51,53,55–58^.

The molecular identities of CDCA components have been a mystery in the PKD field^37,44^. We hypothesize that ciliary ARL13B is a driver in the CDCA pathway. Whereas the complete loss of *Arl13b* causes an ablation of renal cilia, *Arl13b^V358A^* removes ARL13B from cilia while leaving cilia intact, allowing us to directly test the role of ciliary ARL13B in the CDCA pathway to drive renal cystogenesis^48,49,55^. The R79Q mutation in ARL13B ablates ARL13B’s GEF activity for ARL3 *in vitro*^51^. Here we used *Arl13b^R79Q^* mice and demonstrated that ARL13B drives cystogenesis via this critical residue. Lastly, we combined the *Arl13b^V358A^* and *Arl13b^R79Q^* alleles, generating mice in which non-ciliary, cellular ARL13B is enzymatically normal, but ciliary ARL13B has mutant GEF activity. These mutants revealed that the role of ARL13B in cystogenesis is mediated by the residue controlling its ARL3 GEF activity specifically in renal cilia, implicating these ARLs as major drivers of the pro-cystic CDCA pathway.

## Methods

### Mouse strains and procedures

All mice were maintained and cared for in accordance with NIH guidelines and Emory University’s Institutional Animal Care and Use Committee (IACUC). This study was designed and carried out in accordance with ARRIVE guidelines. All informative animals are reported here. Animals were tracked by a unique identifier, and analyses were carried out blinded to genotype of the sample. Animals were fed ad libitum with Purina LabDiet Rodent Diet, Irradiated 5053 and provided free access to water. Cotton nestlets were provided for enrichment and nesting. Animals were monitored daily for signs of dehydration or kidney failure as indicated by hunched posture and weight loss exceeding 10% body weight. All mice were maintained on C57Bl/6J background. *Arl13b* point mutations, including the loss of ciliary ARL13B allele, *Arl13b^V358A^* (MGI:6256969), and loss of the residue controlling the GEF activity of ARL13B for ARL3, *Arl13b^R79Q^* (MGI:6279301), have been previously described^54–56,59^. Conditional alleles include *Pkd1^flox^* (MGI:3612341) and *Arl13b^flox^* (MGI:6107225)^60,61^. Inactivation of conditional floxed alleles was performed using the Pax8^rtTA^ (MGI:3709326) and TetO-Cre (MGI:2679524) transgenes^62,63^. Gene inactivation was achieved by five doses of intraperitoneal (IP) injection of 50 mg/kg doxycycline hyclate in water: three consecutive daily doses administered at 4 weeks, a fourth dose at 6 weeks, and a fifth dose at 8 weeks as recommended by the Polycystic Kidney Disease Research Resource Consortium. Animals were harvested at 18 weeks as described below unless otherwise indicated.

### Animal harvest and tissue processing

Animals were euthanized by isoflurane inhalation and weighed prior to harvest. Blood was collected by cardiac puncture. Following blood collection, the left renal artery was ligated. The left kidney was removed, weighed, and snap frozen for later processing for protein analysis. Animals were perfused with ice cold PBS followed by 4% paraformaldehyde in PBS. After perfusion, the right kidney was removed, weighed, and further fixed in 4% paraformaldehyde for 24 h before embedding in paraffin following standard procedures.

### Blood chemistry

Blood collected via cardiac puncture prior to perfusion was centrifuged at 3500 rpm for 10 min; serum was transferred to a new tube, snap frozen, and stored at -80°C until analysis. Blood urea nitrogen (BUN) and creatinine measurements were carried out by Emory’s Division of Animal Resources (DAR) Diagnostic Services using ACE BUN/Urea and ACE Creatinine Reagents (Alfa Wassermann), respectively.

### Histology

Paraffin embedded kidneys were sectioned at 8-10 µm. For hematoxylin and eosin (H&E) staining, kidney sections were deparaffinized by exposure to SafeClear (Fisher Scientific 23-314629) in 10 min intervals a total of three times, followed by rehydration through a series of decreasing ethanol concentrations for 2 min each (100% ethanol, 100% ethanol, 100% ethanol, 90% ethanol, 70% ethanol, and water). Slides were submerged in Mayer’s hematoxylin (Sigma-Aldrich MHS32) for 7 min, rinsed under running tap water for 15 s, then submerged in 90% ethanol for 30 s. Slides were submerged in eosin (Richard-Allan Scientific 7111) for 30 s, then dehydrated through 100% ethanol four times, and Safeclear a total of three times, then mounted with Cytoseal 60 (Epredia 8310-4). Slides were imaged on an Olympus Nanozoomer slide scanner. Total tissue area was calculated by thresholding in QuPath (v0.6.0)^64^. Cysts were detected by further thresholding on tissue regions with a minimum area of 1500 µm^2^ as previously described^65^. Cystic index was calculated as the sum of the cystic area divided by the total tissue area. For Sirius red/fast green staining, kidney sections were deparaffinized by exposure to SafeClear in 5 min intervals a total of two times, followed by rehydration through a series of decreasing ethanol concentrations for 2 min each (100%, 100%, 95%, 95%, 70%, 50%, water, and water). After rehydration, slides were incubated in 0.04% Fast green FCF (Sigma-Aldrich F7252), 0.1% Direct red 80 (Sigma-Aldrich 365548) in saturated picric aqueous solution (Sigma-Aldrich P6744) for 15 minutes. Slides were dehydrated in 100% ethanol for 5 min, treated with SafeClear for 10 min, then mounted with Cytoseal 60.

### Immunofluorescent Staining

Paraffin-embedded kidney sections were rehydrated as above and antigen retrieval was performed with 10 mM citrate buffer, pH 6.0 (1.26 mM citric acid monohydrate, Fisher Chemical A104 and 8.74 mM sodium citrate dihydrate, Fisher BioReagents BP327). Following antigen retrieval, slides were blocked in 1% heat inactivated goat serum (MP Biomedicals 2939249) and 0.1% Triton X-100 (Fisher BioReagents BP151) in 1x TBS for 1 hr at room temperature. Slides were treated with primary antibodies and incubated overnight at 4°C. Slides were washed three times and incubated with secondary antibodies and Hoechst 33342 (1:5000, Thermo Fisher Scientific H3570, RRID:AB_3675235) for 1 hr at room temperature. Slides were washed three times and mounted with ProLong™ Gold Antifade Mountant (Invitrogen P36930). Primary antibodies used were rabbit anti-SOX9 antibody (1:500, Millipore AB5535, RRID:AB_2239761), rat anti-ARL13B (1:400, BiCell Scientific 90413, RRID:AB_3170226), mouse anti-Acetylated α-Tubulin (1:2000, Sigma-Aldrich T7451, RRID: AB_609894), and rabbit anti-FGFR1OP (1:1000, Proteintech 11343-1-AP, RRID:AB_2103362). Secondary antibodies used were AlexaFluor 488 donkey anti-rat (1:500, Invitrogen A-21208, RRID:AB_2535794), AlexaFluor 568 donkey anti-rabbit (1:500, Thermo Fisher Scientific A10042, RRID:AB_2534017), AlexaFluor 647 goat anti-rabbit (1:500, Jackson ImmunoResearch Labs 111-605-144, RRID:AB_2338078), and AlexaFluor 647 donkey anti-mouse (1:500, Invitrogen A-31573, RRID:AB_2536183). Lotus tetragonolobus lectin (LTL, 1:200, Vector Labs FL-1321) or *Dolichos biflorus* agglutinin (DBA, 1:100, RL-1032) were used as markers of tubular identity. QuPath was used for quantification of SOX9 staining on five or more biological replicates across the entire kidney section in an unbiased manner.

### Cilia Quantification

Z-stack images of immunofluorescent stained kidney sections were captured using an Echo Revolution microscope (voxel depth of 0.82 µm). Cilia length was determined by analyzing Z-stack images with the CiliaQ FIJI Plugin^66^. Cilia were segmented using the acetylated α-tubulin channel, with background subtraction, a rolling ball radius set to 10 pixels, and Triangle thresholding. Post-segmentation analysis was done by filtering cilia with a minimum size of 100 voxels and excluding cilia touching the x or y borders of the images. All other CiliaQ options were left at default settings for segmentation and analysis. Quantification of ARL13B, FGFR1OP, and acetylated α-tubulin was performed using the Cell Counter plugin in FIJI. An average of at least 200 acetylated α-tubulin-positive cilia per animal were counted manually with their respective fluorescent channels across five different fields of view per kidney.

### Western blot

Kidneys were lysed in RIPA Buffer (140 mM NaCl, 10 mM Tris-HCl pH 8.0, 1 mM EDTA, 0.5 mM EGTA, 1% Triton X-100, 0.1% sodium dodecyl sulfate, 0.1% sodium deoxycholate) with Pierce Protease Inhibitor (A32955) and Pierce Phosphatase Inhibitor (88667). Sample concentrations were quantified using Pierce BCA Assay and 40 µg of each sample was loaded onto 10% SDS-PAGE gels. Gels were transferred to 0.2 µm nitrocellulose blotting membranes at 20V for 18 hours. Membranes were blocked in 5% nonfat milk in TBS-T (0.1% Tween-20) for 1 hr at room temperature. Membranes were incubated with mouse anti-HSP90 (1:3000, Santa Cruz Biotechnology sc-13119, RRID:AB_675659) and rabbit anti-α-smooth muscle actin antibody [EPR5368] (1:1000, Abcam ab124964, RRID:AB_11129103), rabbit anti-b-catenin (1:1000, Thermo Fisher Scientific 71-2700, RRID:AB_2533982) or rabbit anti-Cyclin D1 [EPR2241] (1:1500, Abcam ab134175, RRID:AB_2750906) primary antibodies overnight at 4°C. Membranes were washed with TBS-T and incubated with IRDye 680RD donkey anti-mouse (1:10,000, LICORbio 926-68072, RRID:AB_10953628) and IRDye 800CW donkey anti-rabbit (1:10,000, LICORbio 925-32213, RRID:AB_2715510) secondary antibodies for 1 hr at room temperature. Membranes were washed with TBS-T and imaged on an Azure Biosystems imager. Densitometry analysis was performed with ImageLab software version 6.1 (Bio-Rad). Analysis was carried out on three biological replicates.

### Sample Size and Statistics

Sample size and power analysis were calculated using STPLAN (Version 4.5, University of Texas, MD Anderson Cancer Center). For power analysis, normally distributed outcomes and equal variances were assumed. A two-sided test with a significance threshold of 0.05 and 90% power was used. The primary outcome measure used for determining sample size was cystic index as a major indicator of cystic phenotype. A sample size of eight mice would detect a 30% change in mean cystic index of *Arl13b*; *Pkd1^fl/fl^*; *PTCre* mutant mice compared to *Pkd1^fl/fl^*; *PTCre* mice (50%±8.5). Data are presented as mean ± SEM. For kidney morphology and physiology analysis (kidney weight to body weight ratios, cystic indices, blood urea nitrogen, and creatinine), data failed normality testing (Shapiro-Wilk). Data were therefore analyzed with Brown-Forsythe and Welch ANOVA tests, with Dunnett’s T3 correction for multiple comparisons. For western blot and immunofluorescence measurements, data passed normality testing (Shapiro-Wilk), and one-way ANOVA with Tukey’s multiple comparisons test was used for statistical analysis. All p-values are reported in figures compared to Control or *Pkd1^fl/fl^*; PTCre cohorts.

## Results

### Loss of ARL13B from cilia suppressed kidney cyst growth and restored kidney physiology in an adult mouse model of polycystic kidney disease

The CDCA pathway model of PKD pathogenesis posits that its components are in cilia and that loss of critical pathway components suppresses *Pkd1*-dependent cysts. Ciliary proteins are known to play distinct roles in development and adulthood, especially in the kidney^18,67^; here we tested the role of ciliary ARL13B in cystogenesis specifically in an adult model of PKD. To achieve genetic deletion specifically in the kidney epithelium, we used the *Pax8^rtTA^*; *TetO-Cre* system and began doxycycline induction at 4 weeks of age before harvesting animals at 18 weeks (Fig. 1a, see *Methods*). For simplicity, we abbreviate *Pax8^rtTA^*; *TetO-Cre* as *PTCre* throughout this text.

**Figure 1:**
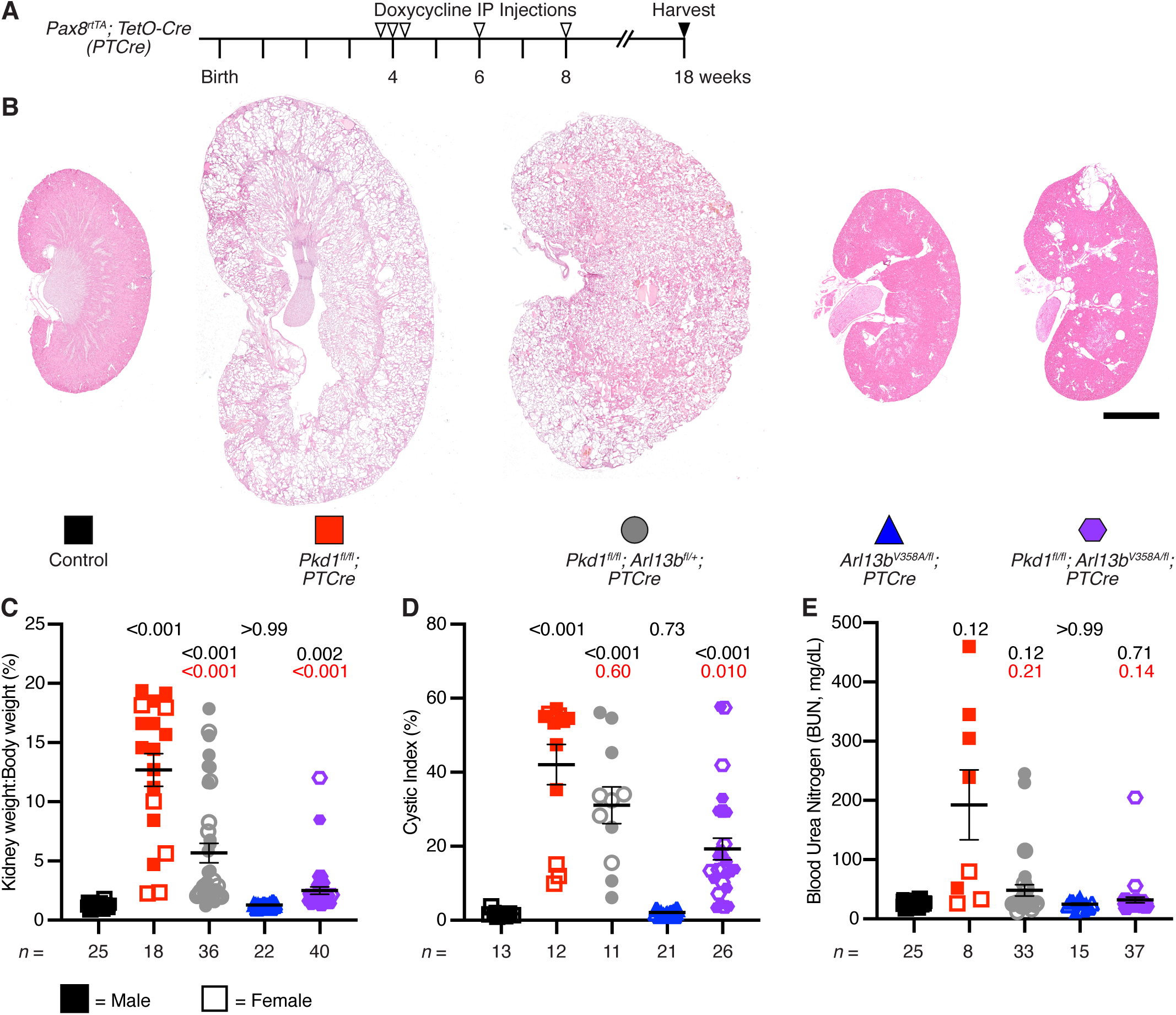
Loss of ciliary ARL13B suppresses severe cystic PC1-deficient phenotypes. Simultaneous loss of ciliary ARL13B and polycystin 1 (PC1, encoded by *Pkd1*) rescued severe cystic phenotypes caused by loss of PC1 alone in an adult *Pkd1*-mouse model. (**a**) Schematic representation of Cre induction and animal harvest experiments. Animals were injected with five doses of doxycycline beginning at 4 weeks of age and harvested at 18 weeks of age. (**b**) Representative hematoxylin and eosin (H&E) staining of kidneys with the indicated genotypes. Scale bar is 2 mm. (**c**) Total kidney weight to body weight ratio. (**d**) Cystic index - cyst area divided by total area of kidney cross section. (**e**) Measurement of blood urea nitrogen (BUN). Symbols in the graphs **c-e** correspond to the genotypes indicated in **b**, with male mice represented by closed symbols and female mice represented by open symbols. Data presented as mean ± SEM; sample sizes (*n*) indicated below each graph. Statistical analysis is Brown-Forsythe and Welch ANOVA tests, with Dunnett’s T3 correction. *P*-values reported for each group compared to control (black text) or *Pkd1^fl/fl^*; *PTCre* (red text) cohorts.

We examined the characteristics of primary cilia across our mutant cohorts, analyzing ciliation, cilia length, and ciliary ARL13B enrichment. To measure ciliation, we stained kidney sections with antibodies to acetylated α-tubulin (ciliary axoneme marker) and FGFR1OP (centrosome marker). We observed similar ciliation rates in all genotypes except for an increase in ciliation in *Pkd1^flox(fl)/fl^*; *PTCre* kidneys (Supplemental Fig. 1a, b). Cilia length increased in *Pkd1^fl/fl^*; *PTCre* kidneys but remained unchanged across all other genotypes (Supplemental Fig. 1a, c). We validated that ARL13B protein was present in control, *Pkd1^fl/fl^*; *PTCre*, and *Pkd1^fl/fl^*; *Arl13b^fl/+^*; *PTCre* kidney cilia and that it was excluded from *Arl13b^V358A/fl^*; *PTCre* and *Pkd1^fl/fl^*; *Arl13b^V358A/fl^*; *PTCre* mutant kidney cilia (Supplemental Fig. 1a, d). Taken together, these results indicate that loss of ciliary ARL13B did not impact cilia frequency or length, demonstrating that these mouse models specifically tested the role of ciliary ARL13B.

As expected, loss of polycystin 1 (PC1, encoded by *Pkd1*) in single mutant *Pkd1^fl/fl^*; *PTCre* animals led to an increase in kidney weight to body weight (KW:BW), cystic index (cyst area divided by total kidney area), blood urea nitrogen (BUN), and creatinine levels compared to control animals (Fig. 1b-e, Supplemental Fig. 2a). We note that the physiological phenotype (increased BUN and creatinine) was significant in male mice (Supplemental Fig. 2b, f) while the gross morphology, including KW:BW and cystic index, were significantly increased for both male and female mice (Supplemental Fig. 2d-i). Since our experimental *Arl13b* mutant alleles were tested over the floxed allele, we next tested whether loss of one copy of *Arl13b* impacted cystic phenotypes. The loss of one copy of *Arl13b* had a mild impact on the *Pkd1* cystic phenotype: *Pkd1^fl/fl^*; *Arl13b^fl/+^*; *PTCre* animals showed a decrease in KW:BW, BUN, and creatinine compared to *Pkd1^fl/fl^*; *PTCre* single mutant animals, while cystic index remained elevated (Fig. 1b-e, Supplemental Fig. 2a). To test whether ciliary ARL13B is a driver of the CDCA pathway, we generated *Pkd1* mice lacking ARL13B specifically in cilia of renal epithelial cells. To do so, we combined the *Arl13b^V358A^* allele, in which ARL13B is undetectable in cilia, with the conditional null *Arl13b^fl^* allele in *Pkd1^fl/fl^* animals. Loss of ciliary ARL13B in single mutant adult *Arl13b^V358A/fl^*; *PTCre* animals did not alter kidney size, morphology, or physiology compared with control animals (Fig. 1b-e, Supplemental Fig. 2a). In contrast, double mutant *Pkd1^fl/fl^*; *Arl13b^V358A/fl^*; *PTCre* animals exhibited a suppression of the severe *Pkd1^fl/fl^*; *PTCre* cystic phenotype, resembling control animals for KW:BW, BUN and creatinine levels (Fig. 1b-c, e, Supplemental Fig. 2a). While cystic index was elevated in *Pkd1^fl/fl^*; *Arl13b^V358A/fl^*; *PTCre* double mutants compared to controls (Fig. 1d), the increase was driven by a few large, focal cysts with most of the tissue exhibiting normal histology (Fig. 1b). These breakthrough cysts did not stain for markers of proximal tubule or collecting duct when stained with lotus tetragonolobus lectin (LTL) or *Dolichos biflorus* agglutinin (DBA), respectively (Supplemental Fig. 3). Overall, our results indicated that ciliary ARL13B directs renal cystogenesis in *Pkd1*-deficient mouse kidneys.

### Loss of ciliary ARL13B suppressed fibrosis in an adult PKD mouse model

Cystogenesis coincides with an accumulation of renal fibrosis, a hallmark of PKD^30,32,33^. As cystic disease progresses, concurrent fibrosis occurs in both mouse models and clinical samples^30,32,33^. We investigated renal fibrosis in our cohorts using Sirius red to stain collagen in fibrotic regions and fast green as a counterstain for non-collagenous regions. We observed Sirius red staining throughout *Pkd1^fl/fl^*; *PTCre* single mutant kidneys (Fig. 2a). The additional loss of one copy of *Arl13b* (*Pkd1^fl/fl^*; *Arl13b^fl/+^*; *PTCre*) did not reduce this phenotype (Fig. 2a). Independent loss of ciliary ARL13B (*Arl13b^V358A/fl^*; *PTCre*) did not result in any observable fibrotic regions (Fig. 2a). *Pkd1^fl/fl^*; *Arl13b^V358A/fl^*; *PTCre* kidneys displayed a marked reduction in fibrosis compared with *Pkd1^fl/fl^*; *PTCre* single mutant kidneys, with most of the kidney free from fibrosis and only small localized fibrotic regions adjacent to cysts (Fig. 2a).

**Figure 2:**
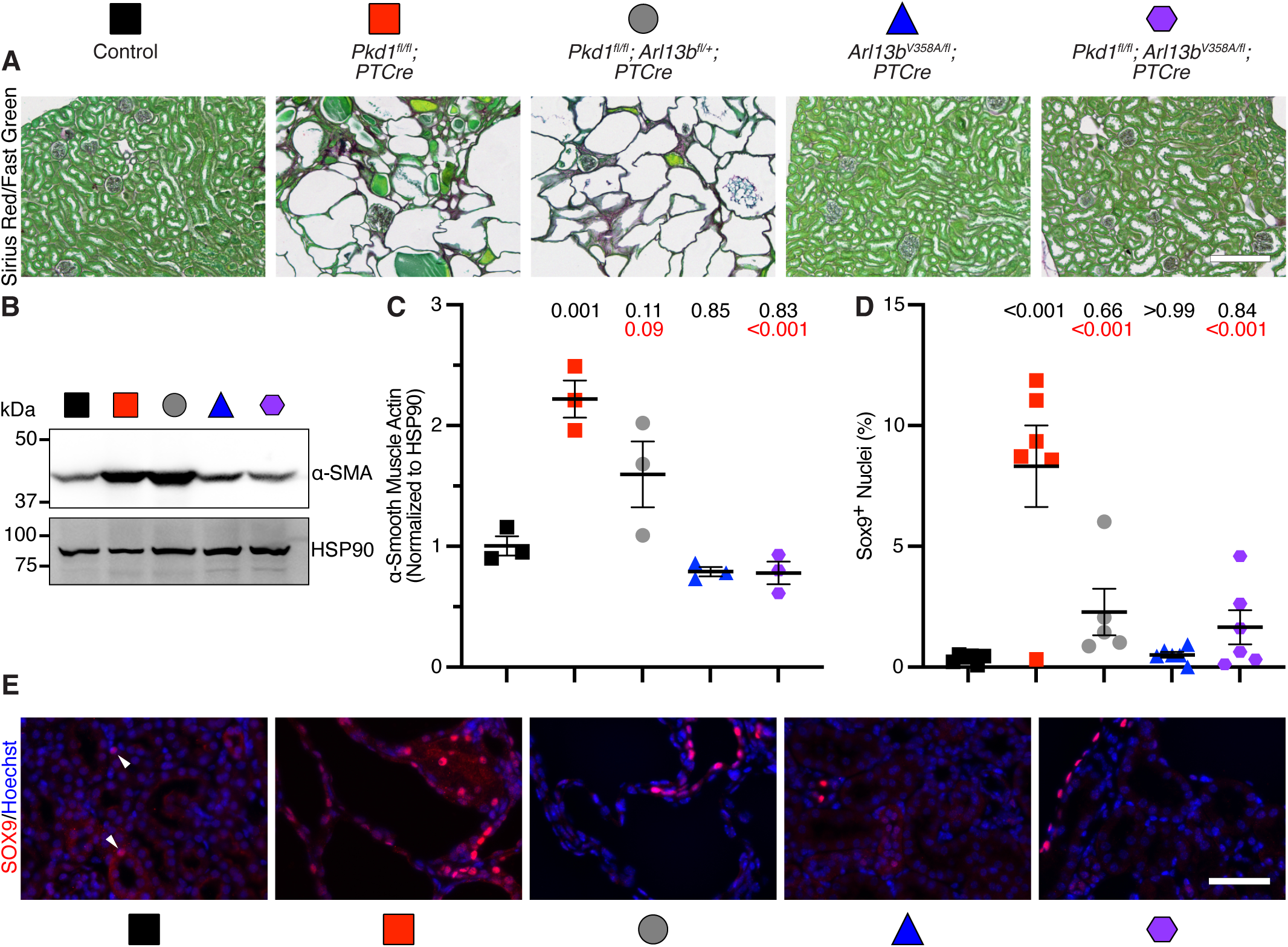
Loss of ciliary ARL13B suppresses fibrosis and injury in adult *Pkd1* mouse models. (**a**) Sirius red and fast green staining reveals fibrosis (red-brown) in cyst-adjacent areas. (**b, c**) Western blot and densitometry analysis of the myofibroblast marker α-smooth muscle actin (α-SMA) in kidney lysates (*n* = 3). (**d, e**) Quantification and images of renal injury marker SOX9 in kidney sections (*n* = 5-6). Scale bars are 200 µm (a) and 50 µm (e). Genotypes in **b-e** correspond to symbol colors and shapes indicated in **a**. Data presented as mean ± SEM. Statistical analysis is one-way ANOVA with Tukey’s multiple comparisons test. *P*-values reported for each group compared to control (black text) or *Pkd1^fl/fl^*; *PTCre* (red text) cohorts.

Renal fibrosis is characterized by elevated levels of α-smooth muscle actin (αSMA), a marker of myofibroblasts^33^. We observed an increase in αSMA protein levels in lysates from *Pkd1^fl/fl^*; *PTCre* single mutant kidneys compared to control kidneys (Fig. 2b, c). The αSMA levels in *Pkd1^fl/fl^*; *Arl13b^fl/+^*; *PTCre* mutant kidney lysates were also elevated, although the levels were not statistically different from *Pkd1^fl/fl^*; *PTCre* single mutant or control kidney lysates (Fig. 2b, c). *Pkd1^fl/fl^*; *Arl13b^V358A/fl^*; *PTCre* double mutant animals displayed similar αSMA levels to control animals, indicating suppression of the fibrotic phenotype observed in *Pkd1^fl/fl^*; *PTCre* single mutants (Fig. 2b, c). While Sirius red staining indicated isolated regions of fibrosis lining the cystic epithelium in our suppression model, the reduction of αSMA levels in kidney lysates indicated that most of the kidney was free from fibrosis. Together, these results indicated that combined loss of PC1 and ciliary exclusion of ARL13B reduced the severe fibrosis seen in loss of PC1 alone.

### Loss of ciliary ARL13B suppressed renal injury in an adult PKD mouse model

In addition to fibrosis, renal injury is a feature of PKD, and SOX9-positive nuclei are indicative of injury in the renal epithelium^25,27,68,69^. We observed a significant increase in SOX9-positive nuclei in *Pkd1^fl/fl^*; *PTCre* single mutant kidneys compared to control kidneys (Fig. 2d,e). Loss of ciliary ARL13B alone (*Arl13b^V358A/fl^*; *PTCre*) showed the same percentage of SOX9-positive nuclei as controls (Fig. 2d,e). Both *Pkd1^fl/fl^*; *Arl13b^V358A/fl^*; *PTCre* double mutant mice and *Pkd1^fl/fl^*; *Arl13b^fl/+^*; *PTCre* mutant mice had SOX9-positive nuclei levels comparable to control animals, indicating suppression of the renal injury observed in PC1 single mutant animals (Fig. 2d, e). Loss of one copy of wild-type *Arl13b* or loss of ciliary ARL13B suppressed the injury observed in *Pkd1*-deleted cystic kidneys. These results indicated that *Pkd1*-dependent renal injury depends on *Arl13b*.

### *Arl13b^R79Q^* suppressed renal cystogenesis in a *Pkd1*-dependent adult mouse model

In addition to its GTPase activity, ARL13B also possesses GEF activity for ARL3, another ARL family GTPase that can localize to cilia^50,51,53,57,58^. Given that loss of ciliary ARL13B suppressed *Pkd1*-dependent cysts (*Pkd1^fl/fl^*; *Arl13b^V358A/fl^*; *PTCre*), we asked whether the residue that controls ARL13B’s GEF activity also suppressed these cysts. We used a mouse expressing ARL13B^R79Q^, a mutant form of ARL13B that cannot activate ARL3 *in vitro*^51,53,56^. We analyzed kidney cilia phenotypes in *Arl13b^R79Q^* mice (*Arl13b^R79Q/fl^*; *PTCre*, *Pkd1^fl/fl^*; *Arl13b^R79Q/fl^*; *PTCre*, *Pkd1^fl/fl^*; *Arl13b^R79Q/R79Q^*; *PTCre* and *Pkd1^fl/fl^*; *Arl13b^R79Q/V358A^*; *PTCre*) and saw no significant change in ciliation, cilia length, or ciliary ARL13B enrichment compared to control tissues (Supplemental Fig. 1). As we observed for the *Arl13b^V358A^* allele, the increase in ciliation in *Pkd1^fl/fl^*; *PTCre* kidneys returned to control levels in all *Arl13b^R79Q^* cohorts (*Pkd1^fl/fl^*; *Arl13b^R79Q/fl^*; *PTCre*, *Pkd1^fl/fl^*; *Arl13b^R79Q/R79Q^*; *PTCre* and *Pkd1^fl/fl^*; *Arl13b^R79Q/V358A^*; *PTCre*; Supplemental Fig. 1a, b). We found cilia length and ciliary ARL13B enrichment from all *Arl13b^R79Q^* cohorts (*Arl13b^R79Q/fl^*; *PTCre*, *Pkd1^fl/fl^*; *Arl13b^R79Q/fl^*; *PTCre*, *Pkd1^fl/fl^*; *Arl13b^R79Q/R79Q^*; *PTCre* and *Pkd1^fl/fl^*; *Arl13b^R79Q/V358A^*; *PTCre*) was comparable to that of control and *Pkd1^fl/fl^*; *PTCre* mutant kidneys (Supplemental Fig. 1). Thus, the *Arl13b^R79Q^* allele isolated the function of the R79Q mutation in ARL13B without affecting its cilia localization.

Mutation of the residue critical for ARL13B GEF activity alone in *Arl13b^R79Q/fl^*; *PTCre* single mutant animals did not result in changes to KW:BW, cystic index, BUN, or creatinine levels compared to control animals (Fig. 3, Supplemental Fig. 4). Simultaneous loss of *Pkd1* with this *Arl13b* mutant (*Pkd1^fl/fl^*; *Arl13b^R79Q/fl^*; *PTCre* and *Pkd1^fl/fl^*; *Arl13b^R79Q/R79Q^*; *PTCre*) suppressed the increased KW:BW, cystic index, BUN, and creatinine levels observed in *Pkd1^fl/fl^*; *PTCre* single mutants (Fig. 3, Supplemental Fig. 4). Together, these results indicated that an ARL13B residue essential for ARL3 GEF activity was required for renal cystogenesis when *Pkd1* was lost.

**Figure 3:**
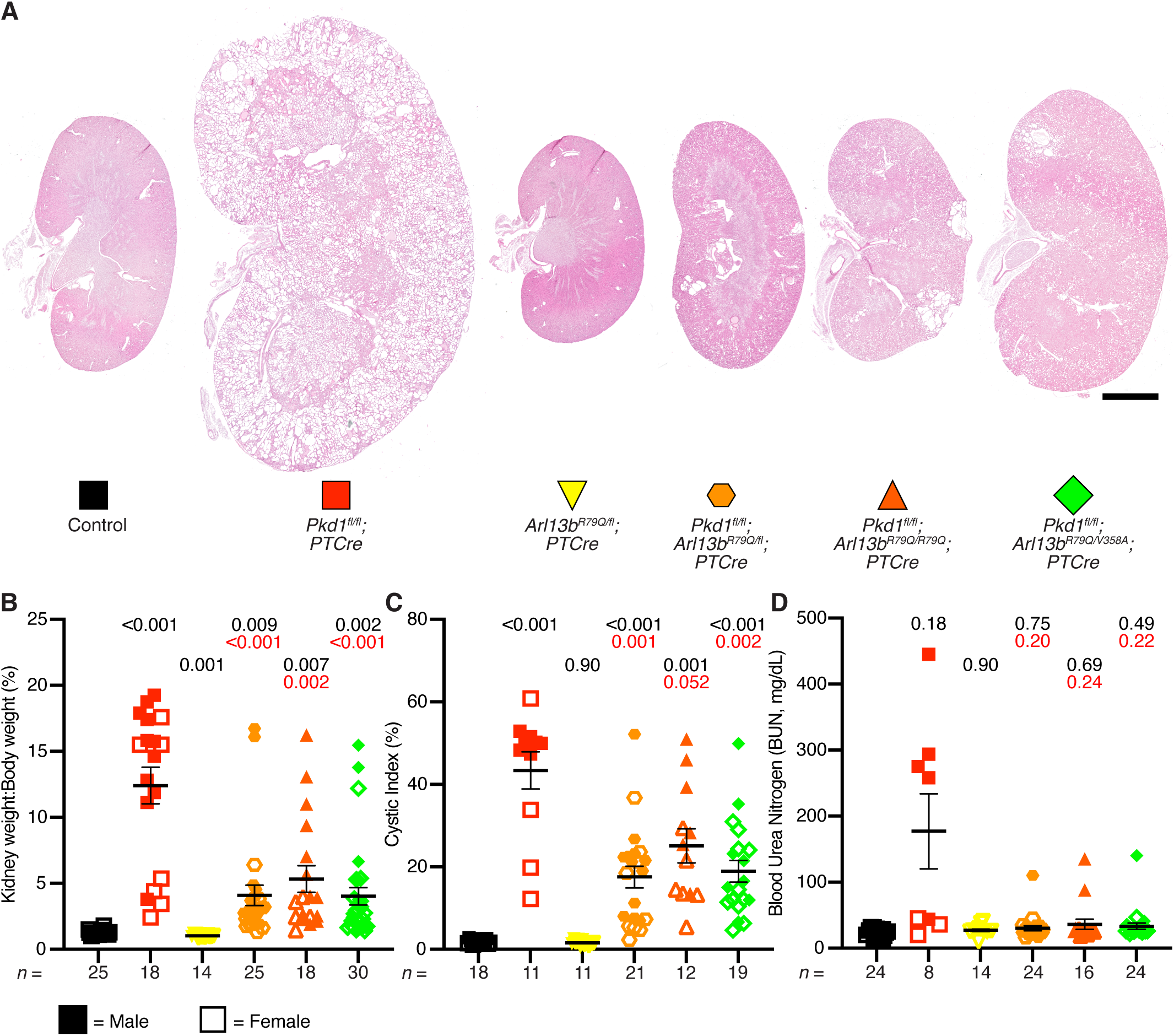
*Arl13b^R79Q^* suppresses severe cystic *Pkd1*-phenotypes. (**a**) Representative H&E staining of kidneys with the indicated genotypes. Scale bar is 2 mm. (**b**) Total kidney weight to body weight ratio. (**c**) Measurement of cystic index. (**d**) Blood urea nitrogen (BUN) measurement. Genotypes in **b-d** correspond to symbol colors and shapes indicated in **a**. Closed symbols indicate male mice and open symbols indicate female mice. Data presented as mean ± SEM; sample sizes indicated below each graph. Statistical analysis is Brown-Forsythe and Welch ANOVA tests, with Dunnett’s T3 correction. *P*-values reported for each group compared to control (black text) or *Pkd1^fl/fl^*; *PTCre* (red text) cohorts.

### *Pkd1*-deficient kidneys expressing *Arl13b^R79Q^* did not exhibit signs of renal fibrosis

Since loss of ciliary ARL13B suppressed fibrosis in *Pkd1* mutant kidneys (Fig. 2a), we asked whether this suppression depended on a residue critical for ARL13B’s GEF activity. We observed levels of Sirius red staining in *Arl13b^R79Q/fl^*; *PTCre* single mutant kidneys comparable to controls (Fig. 4a). As we observed in kidneys lacking ciliary ARL13B (*Pkd1^fl/fl^*; *Arl13b^V358A/fl^*; *PTCre*, Fig. 2a), GEF-deficient ARL13B^R79Q^ expression (*Pkd1^fl/fl^*; *Arl13b^R79Q/R79Q^*; *PTCre* and *Pkd1^fl/fl^*; *Arl13b^R79Q/R79Q^*; *PTCre*) suppressed the severe fibrotic phenotype caused by loss of *Pkd1* alone (Fig. 4a).

**Figure 4:**
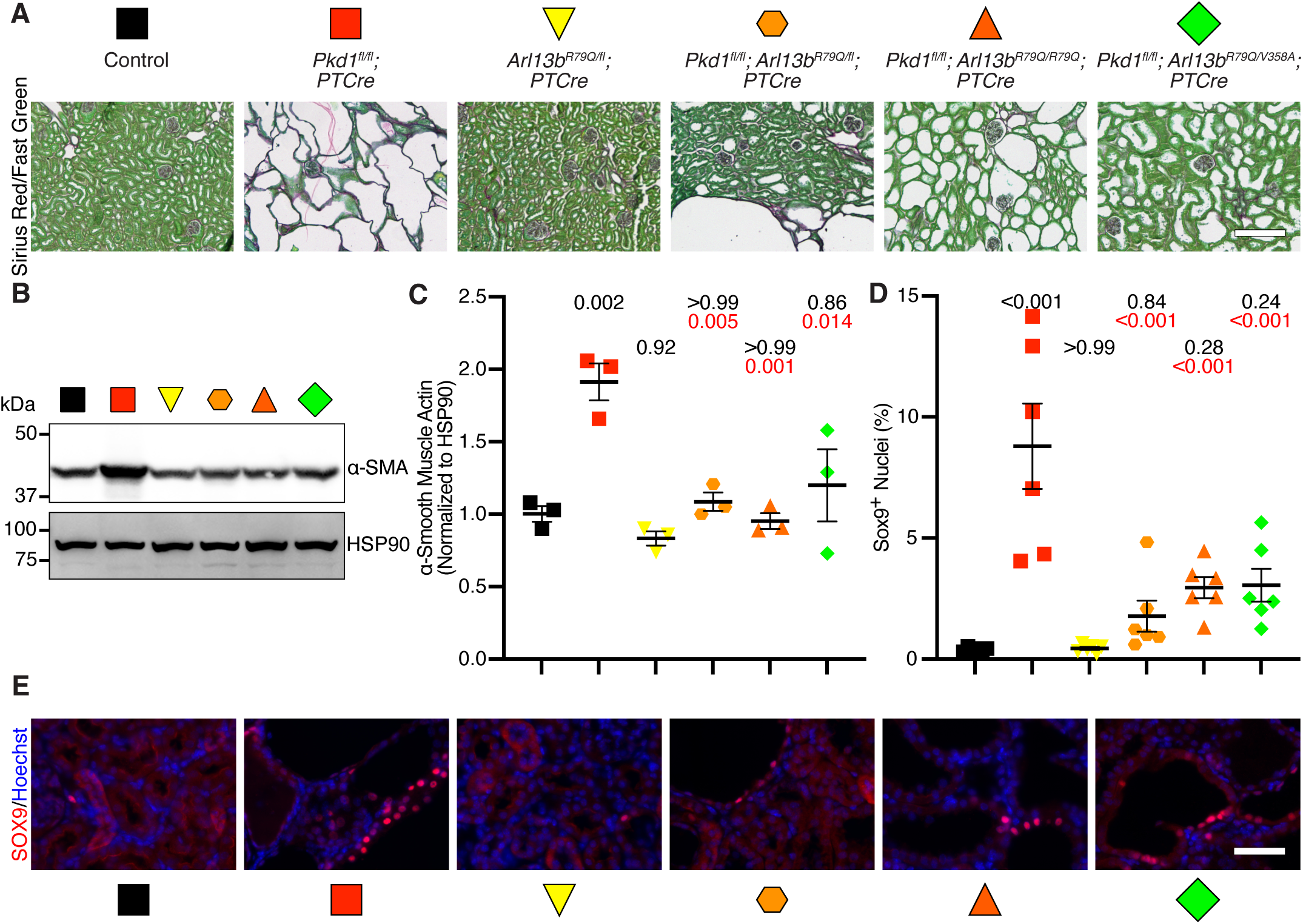
ARL13B^R79Q^ suppresses fibrosis and injury in *Pkd1*-deficient kidneys. (**a**) Sirius red and fast green staining shows fibrosis (red-purple) in cyst-adjacent areas. (**b, c**) Western blot and densitometry analysis of α-SMA in kidney lysates (*n* = 3). (**d, e**) Quantification and images of SOX9 in kidney sections (*n* = 6). Scale bars are 200 µm (**a**) and 50 µm (**e**). Genotypes in **b-e** correspond to symbol colors and shapes indicated in **a**. Data presented as mean ± SEM. Statistical analysis is one-way ANOVA with Tukey’s multiple comparisons test. *P*-values reported for each group compared to control (black text) or *Pkd1^fl/fl^*; *PTCre* (red text) cohorts.

We next looked at αSMA in kidney lysates from these models. As in the previous experiment, single mutant *Pkd1^fl/fl^*; *PTCre* kidney lysates displayed a nearly two-fold increase in αSMA compared to control (Fig. 4b, c; compare to data shown in Fig. 2b, c). Mutation of the residue controlling ARL13B GEF activity in *Arl13b^R79Q/fl^*; *PTCre* mice did not cause any changes in αSMA protein levels compared to control kidney lysates (Fig. 4b,c). *Pkd1^fl/fl^*; *Arl13b^R79Q/fl^*; *PTCre*, and *Pkd1^fl/fl^*; *Arl13b^R79Q/R79Q^*; *PTCre* kidney lysates exhibited levels of αSMA protein comparable to control (Fig. 4b,c). These results indicated that ARL13B^R79Q^ expression suppressed the severe cystic phenotypes and accompanying renal fibrosis caused by loss of *Pkd1* alone.

### Renal injury in a *Pkd1*-dependent mouse model required the residue of ARL13B controlling its ARL3 GEF activity

To investigate renal injury in these mice, we quantified SOX9-positive nuclei. Similar to our previous results, we observed an increase in SOX9-positive nuclei in *Pkd1^fl/fl^*; *PTCre* mutant mice (Fig. 4d, e), indicative of renal epithelial cell injury. Kidney sections from *Arl13b^R79Q/fl^*; *PTCre* mice showed a similar percentage of SOX9-positive nuclei compared to control sections (Fig. 4d, e). The SOX9-positive increase in *Pkd1^fl/fl^*; *PTCre* mutant kidneys was suppressed in *Pkd1^fl/fl^*; *Arl13b^R79Q/fl^*; *PTCre*, and *Pkd1^fl/fl^*; *Arl13b^R79Q/R79Q^*; *PTCre* kidneys, which showed SOX9-positive nuclei numbers comparable to control kidneys (Fig. 4d, e). Taken together, these results indicated that expression of GEF-deficient ARL13B^R79Q^ suppressed the severe cystic phenotypes and associated fibrosis and injury observed when PC1 was lost.

### Adult induced *Pkd1*-deficient kidneys lacking ciliary ARL13B or its GEF activity displayed typical Wnt signaling

To examine the suppression of severe cysts in *Pkd1^fl/fl^*; *PTCre* kidneys by concomitant loss of either ciliary ARL13B or its GEF activity at a molecular level, we examined Wnt signaling as cystic PKD mouse models display elevated Wnt/β-catenin/cyclin D1 signaling^22–24,70^. To assess Wnt signaling in *Pkd1*-dependent cystic kidneys, we immunoblotted kidney lysates for β-catenin and cyclin D1 protein levels (Supplemental Fig. 5a). In agreement with published results, we observed a significant increase in both β-catenin and its transcriptional target cyclin D1 in *Pkd1^fl/fl^*; *PTCre* single mutant kidneys compared to control kidneys (Supplemental Fig. 5a-c). Loss of *Pkd1* and one copy of *Arl13b* (*Pkd1^fl/fl^*; *Arl13b^fl/+^*; *PTCre*) produced β-catenin protein levels comparable to control but cyclin D1 remained elevated (Supplemental Fig. 5a-c). Loss of ciliary ARL13B alone (*Arl13b^V358A/fl^*; *PTCre*) expressed typical β-catenin and cyclin D1 levels compared to control kidney lysates (Supplemental Fig. 5a-c). *Pkd1^fl/fl^*; *Arl13b^V358A/fl^*; *PTCre* double mutant kidneys exhibited β-catenin protein levels comparable to control kidney lysates, indicating a complete reversal of the increase observed in *Pkd1^fl/fl^*; *PTCre* kidney lysates (Supplemental Fig. 5a,b). In contrast, *Pkd1^fl/fl^*; *Arl13b^V358A/fl^*; *PTCre* kidneys exhibited a partial reduction in cyclin D1 levels compared to *Pkd1^fl/fl^*; *PTCre* but not to levels seen in control kidneys (Supplemental Fig. 5a,c).

To examine whether loss of ARL13B GEF activity affected Wnt signaling in an adult *Pkd1* mouse model, we looked at β-catenin and cyclin D1 levels in *Arl13b^R79Q^* animals. Again, *Pkd1^fl/fl^*; *PTCre* kidney lysates exhibited a five- and three-fold increase in β-catenin and cyclin D1 levels compared to control kidney lysates, respectively (Supplemental Fig. 5d-f). *Arl13b^R79Q/fl^*; *PTCre* mice expressed β-catenin and cyclin D1 levels comparable to control animals (Supplemental Fig. 5d-f). *Pkd1^fl/fl^*; *Arl13b^R79Q/fl^*; *PTCre* double mutant mice produced β-catenin and cyclin D1 protein levels comparable to control animals (Supplemental Fig. 5d-f). *Pkd1^fl/fl^*; *Arl13b^R79Q/R79Q^*; *PTCre* kidneys expressed β-catenin typical of control kidneys, but cyclin D1, while reduced, remained significantly elevated compared to control samples (Supplemental Fig. 5d-f). These results indicated that loss of the residue essential for ARL13B GEF activity restored β-catenin and cyclin D1 levels exhibited by *Pkd1* mouse models. Additionally, we observed a dose-dependent phenotype because *Pkd1^fl/fl^*; *Arl13b^R79Q/R79Q^*; *PTCre* kidneys exhibited elevated cyclin D1 protein levels, whereas *Pkd1^fl/fl^*; *Arl13b^R79Q/fl^*; *PTCre* kidneys did not. Taken together, these data indicated that *Pkd1*-deficient kidneys lacking ciliary ARL13B or its GEF activity displayed typical Wnt signaling in an adult PKD mouse model.

### Loss of ARL13B’s GEF activity specifically in cilia suppressed *Pkd1*-dependent phenotypes in an adult mouse model

To explore whether ARL13B drives cystic phenotypes via its GEF activity specifically in cilia, we combined the *Arl13b^R79Q^* and *Arl13b^V358A^* alleles. In *Arl13b^R79Q/V358A^* mice, *Arl13b^R79Q^* expresses ciliary ARL13B lacking the residue controlling GEF activity for ARL3, while *Arl13b^V358A^* codes for non-ciliary (cellular) ARL13B that retains all enzymatic activities. *Pkd1^fl/fl^*; *Arl13b^R79Q/V358A^*; *PTCre* mice exhibited KW:BW, cystic index, BUN, and creatinine levels comparable to control animals, indicating that GEF mutant ARL13B^R79Q^ in cilia suppressed the cysts caused by loss of *Pkd1* alone (Fig. 3, Supplemental Fig. 4). As seen in loss of ARL13B GEF activity in cilia and the cell, GEF mutant ARL13B in cilia and enzymatically normal ARL13B in the cell (*Pkd1^fl/fl^*; *Arl13b^R79Q/V358A^*; *PTCre*) suppressed the severe fibrotic and injury phenotypes caused by loss of *Pkd1* alone (Fig. 4). In summary, *Pkd1^fl/fl^*; *Arl13b^R79Q/V358A^*; *PTCre* mice exhibited diminished renal cystic phenotypes compared to *Pkd1^fl/fl^*; *PTCre* single mutant animals, highlighting a role of ARL13B’s GEF activity specifically in cilia in regulating *Pkd1*-dependent renal cystogenesis (Fig. 5).

**Figure 5.**
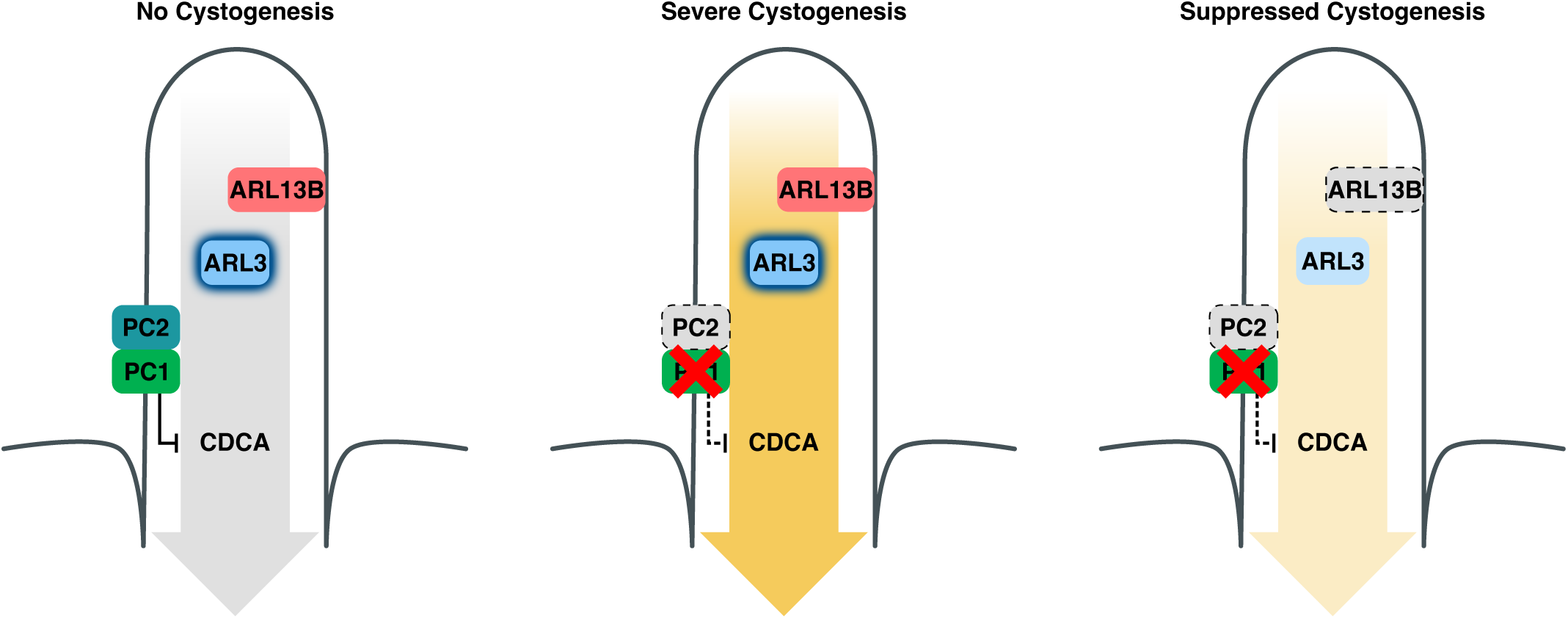
Model showing ciliary ARL13B drives renal cystogenesis in *Pkd1*-deficient kidneys. ARL13B, ARL3, and the polycystin complex PC1/PC2 localize to primary cilia. Under normal conditions, PC1/2 inhibit a pro-cystic CDCA pathway, preventing cystogenesis (**left**). Loss of PC1 (encoded by *Pkd1*) relieves inhibition of the CDCA pathway, resulting in severe cystic phenotypes (**center**). Loss of ciliary ARL13B or mutation of the R79 residue critical for ARL13B’s GEF activity for ARL3 suppresses the severe cystic phenotype caused by loss of PC1 alone (**right**), indicating that ARL13B’s ciliary GEF activity for ARL3 is a major driver of the pro-cystic CDCA pathway.

## Discussion

Researchers have recognized a connection between primary cilia and PKD pathogenesis for several decades^13,16,71^. Results from mouse models strongly suggested the existence of a cilia-dependent cyst activation (CDCA) pathway that drives cystic disease, yet its molecular identity has remained a black box^37,44^. The CDCA pathway hypothesis made two predictions supported by our data: that the CDCA components are in cilia and that loss of function within cilia suppresses polycystin-dependent renal cysts. While this CDCA pathway is one interpretation of suppression in mouse models, it is very difficult to prove this pathway exists until it is properly identified. Our work supports the CDCA model and provides a molecular entry point from which to dissect the CDCA pathway. First, we found that loss of ciliary ARL13B in *Pkd1^fl/fl^*; *Arl13b^V358A/fl^*; *PTCre* double mutants suppressed the severe cystic phenotype seen in *Pkd1^fl/fl^*; *PTCre* single mutant kidneys, indicating ARL13B functions in cilia to drive renal cystogenesis. Second, we showed *Pkd1^fl/fl^*; *Arl13b^R79Q/fl^*; *PTCre* mutants also suppressed the severe cystic phenotype caused by loss of PC1 in *Pkd1^fl/fl^*; *PTCre* kidneys, revealing ARL13B drives renal cystogenesis via the residue controlling its ARL3 GEF activity. Lastly, we observed *Pkd1^fl/fl^*; *Arl13b^R79Q/V358A^*; *PTCre* mutant mice suppressed the severe cystic phenotypes in *Pkd1^fl/fl^*; *PTCre* mutant mice. These results indicated that ARL13B drives renal cystogenesis via the residue controlling its GEF activity for ARL3, specifically in cilia (Fig. 5). Taken together, these data revealed ARL13B to be a driver of the pro-cystic CDCA pathway in the *Pkd1* mouse model. While there are currently no known inhibitors of ARL13B GTPase or GEF activity, nor of ARL3 GTPase activity, our data suggest these enzymatic activities are prime targets for treatment strategies.

We found that loss of *Pkd1* led to an increase in ciliation and cilia length aligning with previous findings^72–74^. Loss of ARL13B’s cilia localization or the residue critical for its ARL3 GEF activity restored ciliation, but did not change its cilia length when compared with control or *Pkd1^fl/fl^* groups. We note that these measurements were performed independent of tubular origin as the renal epithelial cells lining cysts often become de-differentiated and lose their markers of origin identity^74–76^. We showed mouse ARL13B functions in the CDCA pathway via its arginine at amino acid position 79, which is the residue required for its GEF activity *in vitro*^51^. Thus, our data suggest, but do not prove, that ARL13B functions via its GEF activity for ARL3 in the CDCA pathway *in vivo*. While purified mammalian ARL13B^R79Q^ is unable to activate ARL3 *in vitro*, we cannot rule out the possibility of another protein acting as a GEF for ARL3 *in vivo* and have no assay to measure ARL3 activity *in vivo* or specifically within cilia. Future work directly manipulating ARL3 activation state will be needed to directly test the role of ARL3 in mediating suppression of renal cystogenesis. Our results revealed ciliary ARL13B’s GEF activity for ARL3 is a likely mechanism of the CDCA pathway driving cystogenesis.

We note that while most kidney tissue in the genetic suppression models retained normal morphology and physiology, there were isolated “breakthrough” cysts. The most likely explanation for these sporadic cysts is discordant recombination of the floxed alleles as reported for other loci^77,78^; inefficient or delayed recombination of the *Arl13b^fl^* allele may have permitted *Pkd1*-dependent cystogenesis to initiate. Alternatively, the breakthrough cysts could reflect ARL13B cellular GTPase activity playing a minor role in modulating the CDCA pathway in *Pkd1*-mutant cysts. Regardless, we observed grossly normal morphology and renal physiology in *Pkd1*-deficient kidneys, whether cysts were suppressed with *Arl13b^V358A^*, *Arl13b^R79Q^*, or a combination of the two alleles.

Our data are consistent with several previous findings. Loss of the cilia trafficking protein TULP3 suppressed the severe cystic phenotype caused by loss of PC1 alone in adult mouse models of PKD^41^. TULP3 is required for ARL13B ciliary localization in the kidney^41,79^. Our findings suggest that TULP3 mediates cyst suppression via ciliary ARL13B. Additionally, both TULP3 and ARL13B regulate Hedgehog (Hh) signaling, which is implicated in regulating kidney cystogenesis^47,80–83^. While Hh signaling inhibition attenuates cystogenesis in models *in vitro* and *ex vivo*^81–83^, recent work argues that Hh signaling is dispensable for cyst progression *in vivo* in mouse models^84,85^. Despite the misregulation of Hh signaling observed when ARL13B is completely lost, both ARL13B^V358A^ and ARL13B^R79Q^ permit normal Hh signaling^53,54,56^. Thus, our findings are consistent with ARL13B driving cystogenesis independent of its regulation of Hh signaling. Finally, we showed that β-catenin and cyclin D1 are elevated in *Pkd1* mutant kidneys, implicating aberrant Wnt signaling in renal cystogenesis. Loss of ciliary ARL13B or the residue controlling its GEF activity for ARL3 brought β-catenin expression to control levels, but only partially rescued the expression of cyclin D1. Given that either loss of ciliary ARL13B or its GEF activity suppressed the severe cysts in *Pkd1^fl/fl^*; *PTCre* kidneys, we cannot distinguish whether the typical β-catenin and cyclin D1 signaling in those kidneys is due to a role of Wnt in ARL13B–ARL3 signaling axis or because the kidneys in these *Arl13b*; *Pkd1^fl/fl^* double mutants are healthier than *Pkd1^fl/fl^* single mutants overall.

Since the CDCA was first postulated, much work has focused on discovering the molecular identity of a ciliary driver of PKD pathogenesis. Many exciting candidates emerged from these studies, including *Glis2*, *Anks3*, and *Ankmy2*^28,34,42^, however none of the encoded proteins have been found in cilia. GLIS2 and ANKS3 localize to the nucleus and cytoplasm, respectively^28,34^. ANKMY2 regulates the localization of several ciliary proteins, which may be involved in the CDCA pathway; it will be important to investigate this possibility *in vivo*^42,86^. Our work here indicates that ciliary ARL13B is a major component of the CDCA pathway and that this mechanism largely depends on the residue required for ARL13B GEF activity for ARL3. These results provide the field a molecular foothold for discovering additional CDCA components and their mechanisms of action, as well as dissect other ciliary mechanisms driving polycystic kidney disease.

## Acknowledgments

H&E slide scanning in this publication was supported in part by the Cancer Tissue and Pathology Shared Resource of Winship Cancer Institute of Emory University and NIH/NCI under award number P30CA138292. This work was supported by the Emory University Integrated Cellular Imaging Core Facility (RRID:SCR_023534). The *Pax8^rtTA^*, *TetO-Cre*, and *Pkd1^fl/fl^* alleles courtesy the In Vivo PKD Models and Reagents Core of the Polycystic Kidney Disease Research Resource Consortium (PKD RRC). The content is solely the responsibility of the authors and does not necessarily reflect the official views of the National Institutes of Health. Bradley Yoder’s group for helpful discussions and feedback on this project, and the University of Alabama’s Center for Clinical and Translational Science (CCTS) Biostatistics, Epidemiology, & Research Design (BERD) group for statistical guidance. We are grateful specifically to Sarah Bay as well as members of the Caspary lab for feedback, comments, and suggestions in the preparation of this manuscript.

---

**Supplemental Figure 1: Cilia analysis in *Arl13b^V358A^* and *Arl13b^R79Q^* mutant kidneys.** Immunofluorescence analysis reveals ARL13B is not detected in *Arl13b^V358A/V358A^* kidney cilia and is detected in*Arl13b^R79Q/R79Q^* kidney cilia; neither mutation affects ciliation or cilia length. (**a**) Immunofluorescent staining of acetylated α-tubulin (white, AcTub) and ARL13B (green) across various genotypes. (**b**) Quantification of acetylated α-tubulin (AcTub) per FGFR1OP (centrosome marker, FOP). (**c**) Cilia length measurements from Z-stack images. (**d**) Ciliary ARL13B enrichment quantified as ARL13B-positive cilia per AcTub-positive cilia. Genotypes in **b-d** correspond to symbol colors and shapes indicated in **a**. Cilia were analyzed from at least three biological replicates of each genotype (n = 3) and at least 200 cilia were counted from five random fields of view across the kidney. Data presented as mean ± SEM. Statistical analysis is one-way ANOVA with Tukey’s multiple comparisons test. *P*-values reported for each group compared to control (black text) or *Pkd1^fl/fl^*; *PTCre* (red text) cohorts.

**Supplemental Figure 2: Creatinine and sex-specific analyses of loss of ciliary ARL13B and PC1.** Loss of ciliary ARL13B suppressed the elevated creatinine levels observed in *Pkd1*-null kidneys. Morphological phenotypes were present in both male and female mice, while physiological phenotypes were largely driven by male mice. (**a-c**) Serum creatinine analysis across indicated genotypes. (**d-f**) KW:BW, cystic index, and BUN analysis in male mice across indicated genotypes. (**g-i**) KW:BW, cystic index, and BUN analysis in female mice across indicated genotypes. Symbols in the graphs correspond to the genotypes indicated at top of figure, with male mice represented by closed symbols and female mice represented by open symbols. Data presented as mean ± SEM; sample size indicated below each graph. Statistical analysis is Brown-Forsythe and Welch ANOVA tests, with Dunnett’s T3 correction. *P*-values reported for each group compared to control (black text) or *Pkd1^fl/fl^*; *PTCre* (red text) cohorts.

**Supplemental Figure 3: Breakthrough cysts lack tubular markers in *Arl13b* suppression kidneys.** Large breakthrough cysts were observed in the *Arl13b* suppression models of *Pkd1*-dependent cystogenesis. Staining of *Pkd1^fl/fl^*; *Arl13b^V358A/fl^*; *PTCre* mouse kidneys with lotus tetragonolobus lectin (LTL, green) or *Dolichos biflorus* agglutinin (DBA, red) showed these large cysts lacked cellular origin markers.

**Supplemental Figure 4: Creatinine and sex-specific analyses of GEF-deficient ARL13B in *Pkd1*-null kidneys.** Loss of ARL13B GEF activity for ARL3 suppressed the elevated creatinine levels observed in *Pkd1^fl/fl^*; *PTCre* kidneys. Morphological phenotypes were present in both male and female mice, while physiological phenotypes were largely driven by male mice. (**a-c**) Serum creatinine analysis across indicated genotypes. (**d-f**) KW:BW, cystic index, and BUN analysis in male mice across indicated genotypes. (**g-i**) KW:BW, cystic index, and BUN analysis in female mice across indicated genotypes. Symbols in the graphs correspond to the genotypes indicated at top of figure, with male mice represented by closed symbols and female mice represented by open symbols. Data presented as mean ± SEM; sample size indicated below each graph. Statistical analysis is Brown-Forsythe and Welch ANOVA tests, with Dunnett’s T3 correction. *P*-values reported for each group compared to control (black text) or *Pkd1^fl/fl^*; *PTCre* (red text) cohorts.

Supplemental Figure 5: Adult induced *Pkd1*-deficient kidneys lacking ciliary ARL13B or its GEF activity displayed typical Wnt signaling. (**a**) Western blot of WNT signaling components β-catenin and cyclin D1 in kidney lysates. (**b, c**) Densitometry analysis of (**b**) β-catenin and (**c**) cyclin D1 (*n* = 3). (**d**) Western blot of WNT signaling components β-catenin and cyclin D1 in kidney lysates. (**e**, **f**) Densitometry analysis of (**b**) β-catenin and (**c**) cyclin D1 (*n* = 3). Genotypes in **b-c** and **e-f** correspond to symbol colors and shapes indicated in **a** and **d**, respectively. Data presented as mean ± SEM. Statistical analysis is one-way ANOVA with Tukey’s multiple comparisons test. *P*-values reported for each group compared to control (black text) or *Pkd1^fl/fl^*; *PTCre* (red text) cohorts.

**Supplemental Figure 6: Uncropped western blots supporting Figure 2 and Supplemental Figure 5.**

**Supplemental Figure 7: Uncropped western blots supporting Figures 5 and Supplemental Figure 5.**

## References

1. Willey, C., Kamat, S., Stellhorn, R. & Blais, J. Analysis of Nationwide Data to Determine the Incidence and Diagnosed Prevalence of Autosomal Dominant Polycystic Kidney Disease in the USA: 2013–2015. Kidney Diseases 5, 107–117 (2019).

2. Willey, C.J. et al. Prevalence of autosomal dominant polycystic kidney disease in the European Union. Nephrology Dialysis Transplantation 32, 1356–1363 (2016).

3. Bergmann, C. et al. Polycystic kidney disease. Nat Rev Dis Primers 4, 50 (2018).

4. Lanktree, M.B. et al. Prevalence Estimates of Polycystic Kidney and Liver Disease by Population Sequencing. Journal of the American Society of Nephrology 29, 2593–2600 (2018).

5. Lanktree, M.B., Haghighi, A., di Bari, I., Song, X. & Pei, Y. Insights into Autosomal Dominant Polycystic Kidney Disease from Genetic Studies. *Clinical Journal of the American Society of Nephrology*, CJN.02320220 (2020).

6. Cornec-Le Gall, E., Alam, A. & Perrone, R.D. Autosomal dominant polycystic kidney disease. Lancet 393, 919–935 (2019).

7. Cornec-Le Gall, E., Torres, V.E. & Harris, P.C. Genetic Complexity of Autosomal Dominant Polycystic Kidney and Liver Diseases. J Am Soc Nephrol 29, 13–23 (2018).

8. Hughes, J. et al. The polycystic kidney disease 1 (PKD1) gene encodes a novel protein with multiple cell recognition domains. Nat Genet 10, 151–60 (1995).

9. Mochizuki, T. et al. PKD2, a gene for polycystic kidney disease that encodes an integral membrane protein. Science 272, 1339–42 (1996).

10. The International Polycystic Kidney Disease Consortium. Polycystic kidney disease: the complete structure of the PKD1 gene and its protein. The International Polycystic Kidney Disease Consortium. Cell 81, 289–98 (1995).

11. Huangfu, D. et al. Hedgehog signalling in the mouse requires intraflagellar transport proteins. Nature 426, 83–7 (2003).

12. Barr, M.M. & Sternberg, P.W. A polycystic kidney-disease gene homologue required for male mating behaviour in C. elegans. Nature 401, 386–9 (1999).

13. Pazour, G.J. et al. Chlamydomonas IFT88 and its mouse homologue, polycystic kidney disease gene tg737, are required for assembly of cilia and flagella. J Cell Biol 151, 709–18 (2000).

14. Anvarian, Z., Mykytyn, K., Mukhopadhyay, S., Pedersen, L.B. & Christensen, S.T. Cellular signalling by primary cilia in development, organ function and disease. Nature Reviews Nephrology 15, 199–219 (2019).

15. Garcia-Gonzalo, F.R. et al. Phosphoinositides Regulate Ciliary Protein Trafficking to Modulate Hedgehog Signaling. Dev Cell 34, 400–409 (2015).

16. Yoder, B.K., Hou, X. & Guay-Woodford, L.M. The polycystic kidney disease proteins, polycystin-1, polycystin-2, polaris, and cystin, are co-localized in renal cilia. J Am Soc Nephrol 13, 2508–16 (2002).

17. Andrews, P.M. & Porter, K.R. A scanning electron microscopic study of the nephron. Am J Anat 140, 81–115 (1974).

18. Davenport, J.R. et al. Disruption of intraflagellar transport in adult mice leads to obesity and slow-onset cystic kidney disease. Curr Biol 17, 1586–94 (2007).

19. Zhang, Q., Taulman, P.D. & Yoder, B.K. Cystic Kidney Diseases: All Roads Lead to the Cilium. Physiology 19, 225–230 (2004).

20. Walker, R.V. et al. Ciliary exclusion of Polycystin-2 promotes kidney cystogenesis in an autosomal dominant polycystic kidney disease model. Nat Commun 10, 4072 (2019).

21. Franco, I. et al. Phosphoinositide 3-Kinase-C2alpha Regulates Polycystin-2 Ciliary Entry and Protects against Kidney Cyst Formation. J Am Soc Nephrol 27, 1135–44 (2016).

22. Kim, S. et al. The polycystin complex mediates Wnt/Ca2+ signalling. Nature Cell Biology 18, 752–764 (2016).

23. Conduit, S.E. et al. β-catenin ablation exacerbates polycystic kidney disease progression. Human Molecular Genetics 28, 230–244 (2018).

24. Lee, E.J. et al. TAZ/Wnt-β-catenin/c-MYC axis regulates cystogenesis in polycystic kidney disease. Proceedings of the National Academy of Sciences 117, 29001–29012 (2020).

25. Leonhard, W.N. et al. Scattered Deletion of PKD1 in Kidneys Causes a Cystic Snowball Effect and Recapitulates Polycystic Kidney Disease. Journal of the American Society of Nephrology 26, 1322–1333 (2015).

26. Wang, Q. et al. Adenylyl cyclase 5 deficiency reduces renal cyclic AMP and cyst growth in an orthologous mouse model of polycystic kidney disease. Kidney International 93, 403–415 (2018).

27. Zimmerman, K.A. et al. Tissue-Resident Macrophages Promote Renal Cystic Disease. J Am Soc Nephrol (2019).

28. Wei, Z. et al. Anks3 mediates cilia dependent polycystin signaling and is essential for adult kidney homeostasis. bioRxiv, 2025.04.22.649832 (2025).

29. Chang, M.Y. et al. Haploinsufficiency of Pkd2 is associated with increased tubular cell proliferation and interstitial fibrosis in two murine Pkd2 models. Nephrology Dialysis Transplantation 21, 2078–2084 (2006).

30. Song, C.J., Zimmerman, K.A., Henke, S.J. & Yoder, B.K. Inflammation and Fibrosis in Polycystic Kidney Disease. in Kidney Development and Disease (ed. Miller, R.K.) 323–344 (Springer International Publishing, Cham, 2017).

31. Dong, K. et al. Renal plasticity revealed through reversal of polycystic kidney disease in mice. Nature Genetics 53, 1649–1663 (2021).

32. Lantinga-van Leeuwen, I.S., et al. Lowering of Pkd1 expression is sufficient to cause polycystic kidney disease. Hum Mol Genet 13, 3069–77 (2004).

33. Jiang, S.-T. et al. Defining a Link with Autosomal-Dominant Polycystic Kidney Disease in Mice with Congenitally Low Expression of *Pkd1*. The American Journal of Pathology 168, 205–220 (2006).

34. Zhang, C. et al. Glis2 is an early effector of polycystin signaling and a target for therapy in polycystic kidney disease. Nature Communications 15, 3698 (2024).

35. He, J. et al. Inhibiting Focal Adhesion Kinase Ameliorates Cyst Development in Polycystin-1–Deficient Polycystic Kidney Disease in Animal Model. Journal of the American Society of Nephrology 32, 2159–2174 (2021).

36. Lee, K., Boctor, S., Barisoni, L.M.C. & Gusella, G.L. Inactivation of Integrin-β1 Prevents the Development of Polycystic Kidney Disease after the Loss of Polycystin-1. Journal of the American Society of Nephrology 26, 888–895 (2015).

37. Ma, M., Tian, X., Igarashi, P., Pazour, G.J. & Somlo, S. Loss of cilia suppresses cyst growth in genetic models of autosomal dominant polycystic kidney disease. Nat Genet 45, 1004–12 (2013).

38. Zhang, C. et al. Cyclin-Dependent Kinase 1 Activity Is a Driver of Cyst Growth in Polycystic Kidney Disease. Journal of the American Society of Nephrology, ASN.2020040511 (2020).

39. Kraus, A. et al. P2Y2R and Cyst Growth in Polycystic Kidney Disease. Journal of the American Society of Nephrology, 10.1681/ASN.0000000000000416 (2024).

40. Cabrita, I. et al. Cyst growth in ADPKD is prevented by pharmacological and genetic inhibition of TMEM16A in vivo. Nature Communications 11, 4320 (2020).

41. Legue, E. & Liem, K.F., Jr. Tulp3 Is a Ciliary Trafficking Gene that Regulates Polycystic Kidney Disease. Curr Biol 29, 803–812 e5 (2019).

42. Hwang, S.-h., et al. Kidney cystogenesis in embryonic- and adult-onset ADPKD is suppressed from lack of adenylyl cyclase targeting to cilia. bioRxiv, 2025.05.27.656198 (2025).

43. Walker, R.V. et al. Cilia-Localized Counterregulatory Signals as Drivers of Renal Cystogenesis. Front Mol Biosci 9, 936070 (2022).

44. Ma, M., Gallagher, A.R. & Somlo, S. Ciliary Mechanisms of Cyst Formation in Polycystic Kidney Disease. Cold Spring Harb Perspect Biol 9(2017).

45. Antignac, C. et al. The Future of Polycystic Kidney Disease Research--As Seen By the 12 Kaplan Awardees. J Am Soc Nephrol 26, 2081–95 (2015).

46. Sun, Z. et al. A genetic screen in zebrafish identifies cilia genes as a principal cause of cystic kidney. Development 131, 4085–93 (2004).

47. Caspary, T., Larkins, C.E. & Anderson, K.V. The graded response to Sonic Hedgehog depends on cilia architecture. Dev Cell 12, 767–78 (2007).

48. Li, Y. et al. Deletion of ADP Ribosylation Factor-Like GTPase 13B Leads to Kidney Cysts. J Am Soc Nephrol 27, 3628–3638 (2016).

49. Seixas, C. et al. Arl13b and the exocyst interact synergistically in ciliogenesis. Mol Biol Cell 27, 308–20 (2016).

50. Gotthardt, K. et al. A G-protein activation cascade from Arl13B to Arl3 and implications for ciliary targeting of lipidated proteins. Elife 4(2015).

51. Ivanova, A.A. et al. Biochemical characterization of purified mammalian ARL13B protein indicates that it is an atypical GTPase and ARL3 guanine nucleotide exchange factor (GEF). J Biol Chem 292, 11091–11108 (2017).

52. Higginbotham, H. et al. Arl13b in primary cilia regulates the migration and placement of interneurons in the developing cerebral cortex. Dev Cell 23, 925–38 (2012).

53. Mariani, L.E. et al. Arl13b regulates Shh signaling from both inside and outside the cilium. Molecular Biology of the Cell 27, 3780–3790 (2016).

54. Gigante, E.D., Taylor, M.R., Ivanova, A.A., Kahn, R.A. & Caspary, T. ARL13B regulates Sonic hedgehog signaling from outside primary cilia. eLife 9, e50434 (2020).

55. Van Sciver, R.E., Long, A.B., Katz, H.G., Gigante, E.D. & Caspary, T. Ciliary ARL13B inhibits developmental kidney cystogenesis in mouse. Dev Biol 500, 1–9 (2023).

56. Suciu, S.K., Long, A.B. & Caspary, T. Smoothened and ARL13B are critical in mouse for superior cerebellar peduncle targeting. Genetics 218(2021).

57. Cantagrel, V. et al. Mutations in the cilia gene ARL13B lead to the classical form of Joubert syndrome. Am J Hum Genet 83, 170–9 (2008).

58. Humbert, M.C. et al. ARL13B, PDE6D, and CEP164 form a functional network for INPP5E ciliary targeting. Proc Natl Acad Sci U S A 109, 19691–6 (2012).

59. Ferent, J. et al. The Ciliary Protein Arl13b Functions Outside of the Primary Cilium in Shh-Mediated Axon Guidance. Cell Reports 29, 3356–3366.e3 (2019).

60. Piontek, K.B. et al. A functional floxed allele of Pkd1 that can be conditionally inactivated in vivo. J Am Soc Nephrol 15, 3035–43 (2004).

61. Su, C.Y., Bay, S.N., Mariani, L.E., Hillman, M.J. & Caspary, T. Temporal deletion of Arl13b reveals that a mispatterned neural tube corrects cell fate over time. Development 139, 4062–71 (2012).

62. Traykova-Brauch, M. et al. An efficient and versatile system for acute and chronic modulation of renal tubular function in transgenic mice. Nat Med 14, 979–84 (2008).

63. Perl, A.K., Wert, S.E., Nagy, A., Lobe, C.G. & Whitsett, J.A. Early restriction of peripheral and proximal cell lineages during formation of the lung. Proc Natl Acad Sci U S A 99, 10482–7 (2002).

64. Bankhead, P. et al. QuPath: Open source software for digital pathology image analysis. Scientific Reports 7, 16878 (2017).

65. Wilson, L. et al. Chronic activation of AMP-activated protein kinase leads to early-onset polycystic kidney phenotype. Clinical Science 135, 2393–2408 (2021).

66. Hansen, J.N., Rassmann, S., Stüven, B., Jurisch-Yaksi, N. & Wachten, D. CiliaQ: a simple, open-source software for automated quantification of ciliary morphology and fluorescence in 2D, 3D, and 4D images. The European Physical Journal E 44, 18 (2021).

67. Piontek, K., Menezes, L.F., Garcia-Gonzalez, M.A., Huso, D.L. & Germino, G.G. A critical developmental switch defines the kinetics of kidney cyst formation after loss of Pkd1. Nat Med 13, 1490–5 (2007).

68. Aggarwal, S. et al. SOX9 switch links regeneration to fibrosis at the single-cell level in mammalian kidneys. Science 383, eadd6371 (2024).

69. Seeger-Nukpezah, T., Geynisman, D.M., Nikonova, A.S., Benzing, T. & Golemis, E.A. The hallmarks of cancer: relevance to the pathogenesis of polycystic kidney disease. Nat Rev Nephrol 11, 515–34 (2015).

70. Happé, H. et al. Toxic tubular injury in kidneys from Pkd1-deletion mice accelerates cystogenesis accompanied by dysregulated planar cell polarity and canonical Wnt signaling pathways. Human Molecular Genetics 18, 2532–2542 (2009).

71. Pazour, G.J., San Agustin, J.T., Follit, J.A., Rosenbaum, J.L. & Witman, G.B. Polycystin-2 localizes to kidney cilia and the ciliary level is elevated in orpk mice with polycystic kidney disease. Curr Biol 12, R378–80 (2002).

72. Liu, J. et al. Non-parallel recombination limits cre-loxP-based reporters as precise indicators of conditional genetic manipulation. genesis 51, 436–442 (2013).

73. Vooijs, M., Jonkers, J. & Berns, A. A highly efficient ligand-regulated Cre recombinase mouse line shows that *LoxP* recombination is position dependent. EMBO reports 2, 292–297 (2001).

74. Hwang, S.H. et al. Tulp3 Regulates Renal Cystogenesis by Trafficking of Cystoproteins to Cilia. Curr Biol 29, 790–802 e5 (2019).

75. Mukhopadhyay, S. et al. The ciliary G-protein-coupled receptor Gpr161 negatively regulates the Sonic hedgehog pathway via cAMP signaling. Cell 152, 210–23 (2013).

76. Silva, L.M. et al. Inhibition of Hedgehog signaling suppresses proliferation and microcyst formation of human Autosomal Dominant Polycystic Kidney Disease cells. Sci Rep 8, 4985 (2018).

77. Hsieh, C.-L., Jerman, S.J. & Sun, Z. Non-cell-autonomous activation of hedgehog signaling contributes to disease progression in a mouse model of renal cystic ciliopathy. Human Molecular Genetics 31, 4228–4240 (2022).

78. Tran, P.V. et al. Downregulating hedgehog signaling reduces renal cystogenic potential of mouse models. J Am Soc Nephrol 25, 2201–12 (2014).

79. Ma, M., Legué, E., Tian, X., Somlo, S. & Liem, K.F.J. Cell-Autonomous Hedgehog Signaling Is Not Required for Cyst Formation in Autosomal Dominant Polycystic Kidney Disease. Journal of the American Society of Nephrology 30, 2103–2111 (2019).

80. Gombart, S., Houghtaling, S., Ho, T.-H. & Beier, D.R. Inhibition of Hedgehog signaling does not mitigate polycystic kidney disease severity in a Pkd1 mutant mouse model. Journal of Cell Science, jcs.264133 (2025).

81. Somatilaka, B.N. et al. Ankmy2 Prevents Smoothened-Independent Hyperactivation of the Hedgehog Pathway via Cilia-Regulated Adenylyl Cyclase Signaling. Developmental Cell 54, 710–726.e8 (2020).

