## Supplemental Figures for "Ciliary ARL13B drives renal cystogenesis via its GEF activity"

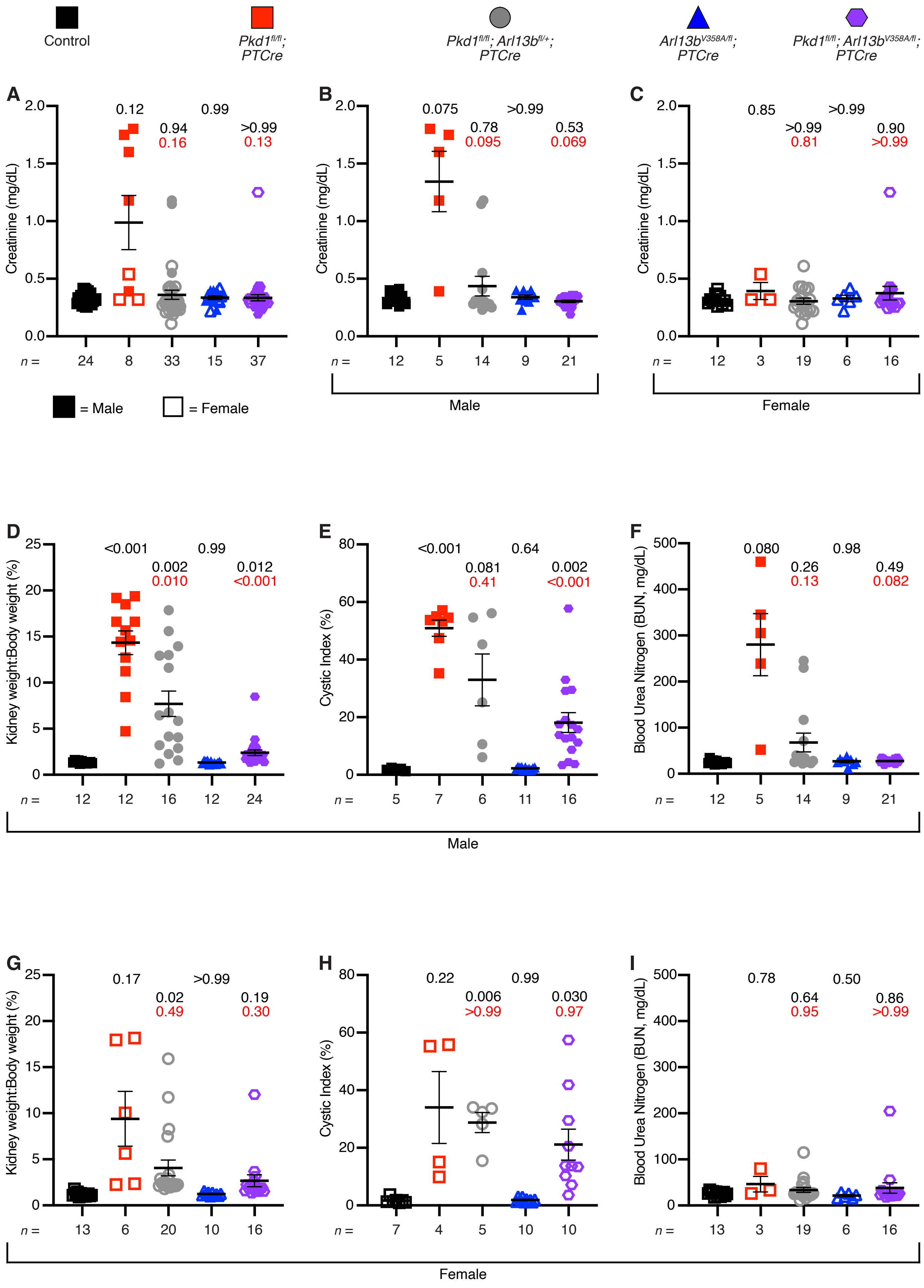

Control
Pkd1fl/fl; PTCre
Pkd1fl/fl; Arl13bfl/+; PTCre
Arl13bV358A/fl; PTCre
Pkd1fl/fl; Arl13bV358A/fl; PTCre
A
B
C
Creatinine (mg/dL)
0.12
0.94
0.16
0.99
>0.99
0.13
n = 24 8 33 15 37
= Male
= Female
0.075
0.78
0.095
>0.99
0.53
0.069
n = 12 5 14 9 21
Male
0.85
>0.99
0.81
>0.99
0.90
>0.99
n = 12 3 19 6 16
Female
D
E
F
Kidney weight:Body weight (%)
<0.001
0.002
0.010
0.99
0.012
<0.001
n = 12 12 16 12 24
Cystic Index (%)
<0.001
0.081
0.41
0.64
0.002
<0.001
n = 5 7 6 11 16
Blood Urea Nitrogen (BUN, mg/dL)
0.080
0.26
0.13
0.98
0.49
0.082
n = 12 5 14 9 21
Male
G
H
I
Kidney weight:Body weight (%)
0.17
0.02
0.49
>0.99
0.19
0.30
n = 13 6 20 10 16
Cystic Index (%)
0.22
0.006
>0.99
0.99
0.030
0.97
n = 7 4 5 10 10
Blood Urea Nitrogen (BUN, mg/dL)
0.78
0.64
0.95
0.50
0.86
>0.99
n = 13 3 19 6 16
Female

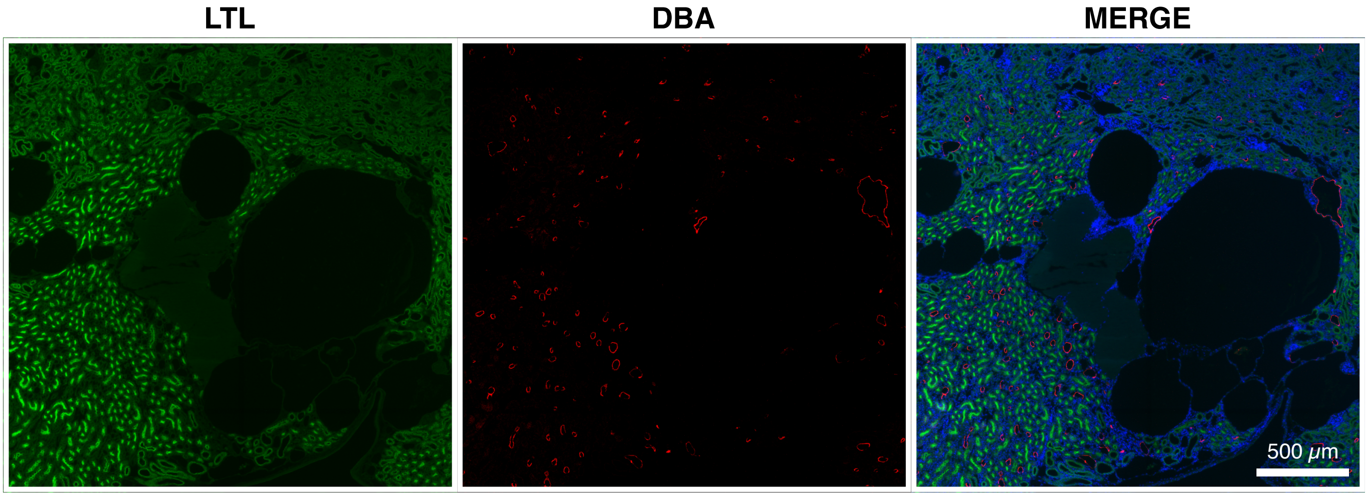

LTL
DBA
MERGE
500 μm

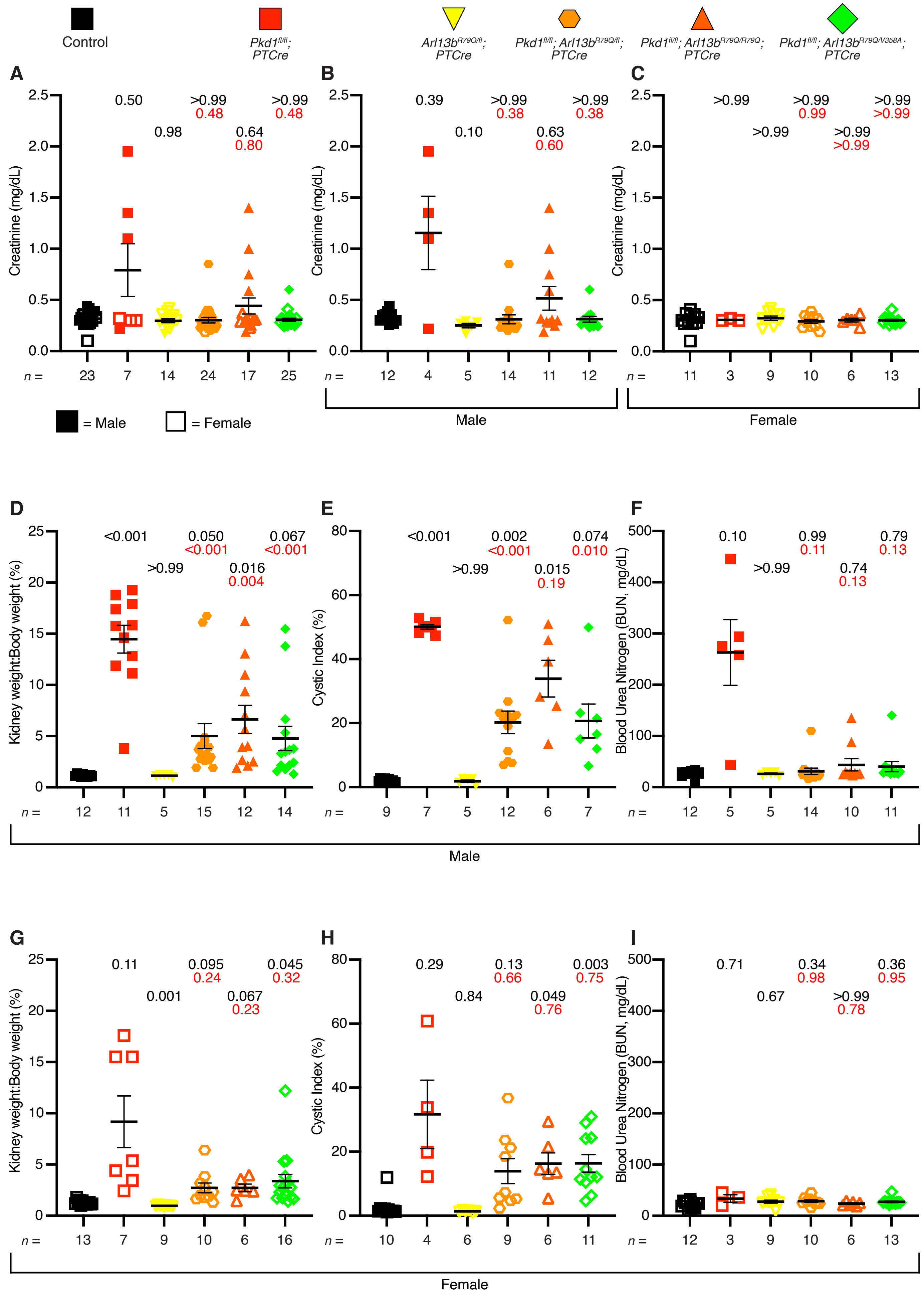
Control
Pkd1fl/fl; PTCre
Arl13bR79Q/fl; PTCre
Pkd1fl/fl; Arl13bR79Q/fl; PTCre
Pkd1fl/fl; Arl13bR79Q/R79Q; PTCre
Pkd1fl/fl; Arl13bR79Q/V358A; PTCre
A
B
C
Creatinine (mg/dL)
0.50
0.98
>0.99
0.48
0.64
0.80
>0.99
0.48
n = 23 7 14 24 17 25
0.39
0.10
>0.99
0.38
0.63
0.60
>0.99
0.38
n = 12 4 5 14 11 12
>0.99
>0.99
>0.99
0.99
>0.99
>0.99
>0.99
>0.99
n = 11 3 9 10 6 13
= Male
= Female
Male
Female
D
E
F
Kidney weight:Body weight (%)
<0.001
>0.99
0.050
<0.001
0.016
0.004
0.067
<0.001
n = 12 11 5 15 12 14
Cystic Index (%)
<0.001
>0.99
0.002
<0.001
0.015
0.19
0.074
0.010
n = 9 7 5 12 6 7
Blood Urea Nitrogen (BUN, mg/dL)
0.10
>0.99
0.99
0.11
0.74
0.13
0.79
0.13
n = 12 5 5 14 10 11
Male
G
H
I
Kidney weight:Body weight (%)
0.11
0.001
0.095
0.24
0.067
0.23
0.045
0.32
n = 13 7 9 10 6 16
Cystic Index (%)
0.29
0.84
0.13
0.66
0.049
0.76
0.003
0.75
n = 10 4 6 9 6 11
Blood Urea Nitrogen (BUN, mg/dL)
0.71
0.67
0.34
0.98
>0.99
0.78
0.36
0.95
n = 12 3 9 10 6 13
Female

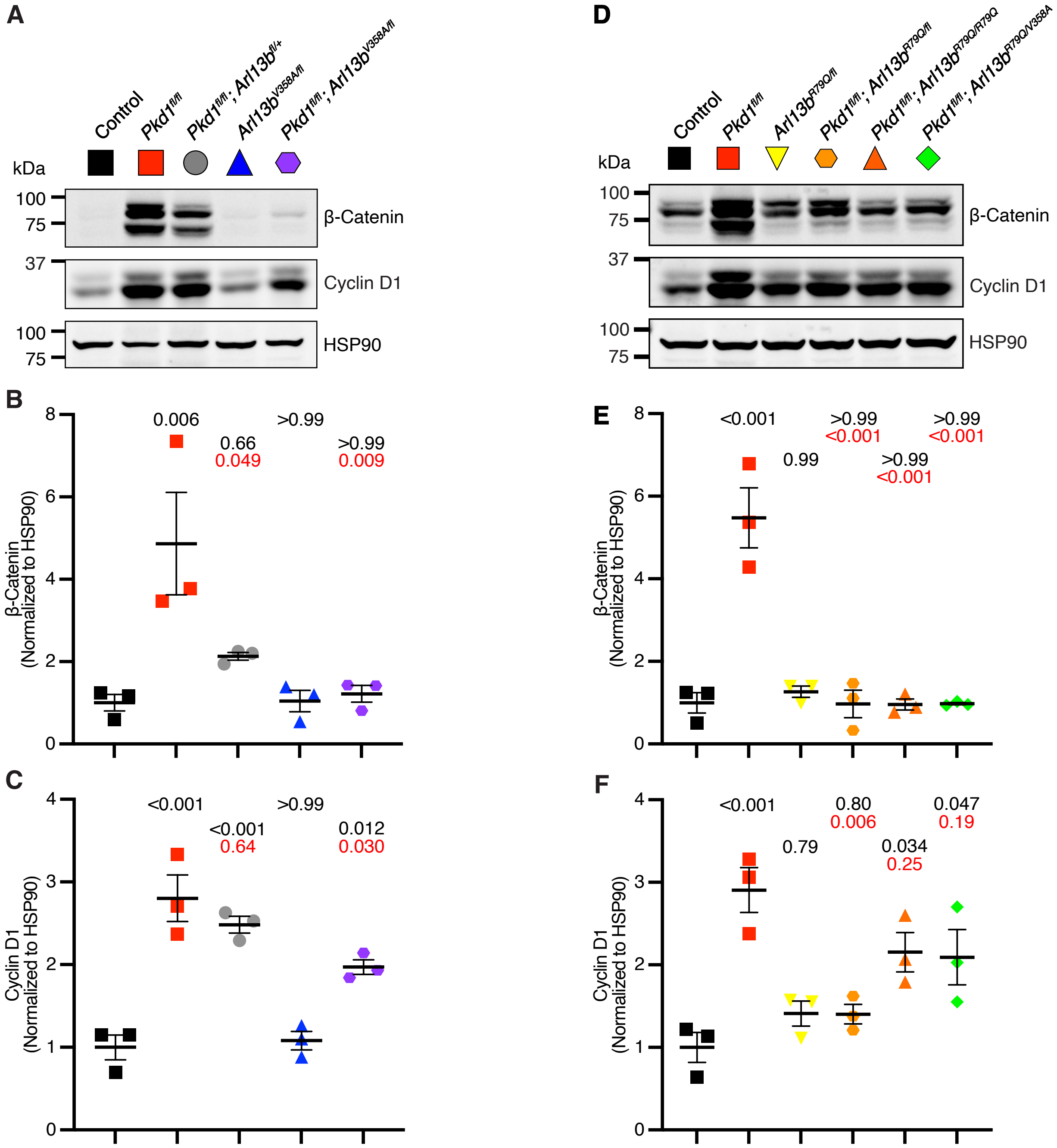
A
Control
Pkd1fl/fl
Pkd1fl/fl; Arl13bfl/+
Arl13bV358A/fl
Pkd1fl/fl; Arl13bV358A/fl
kDa
100
75
β-Catenin
37
Cyclin D1
100
75
HSP90
B
β-Catenin (Normalized to HSP90)
0.006
0.66
0.049
>0.99
>0.99
0.009
C
Cyclin D1 (Normalized to HSP90)
<0.001
<0.001
0.64
>0.99
0.012
0.030
D
Control
Pkd1fl/fl
Arl13bR79Q/fl
Pkd1fl/fl; Arl13bR79Q/fl
Pkd1fl/fl; Arl13bR79Q/R79Q
Pkd1fl/fl; Arl13bR79Q/V358A
kDa
100
75
β-Catenin
37
Cyclin D1
100
75
HSP90
E
β-Catenin (Normalized to HSP90)
<0.001
0.99
>0.99
<0.001
>0.99
<0.001
>0.99
<0.001
F
Cyclin D1 (Normalized to HSP90)
<0.001
0.79
0.80
0.006
0.034
0.25
0.047
0.19

Full-length western blots from Figure 2b

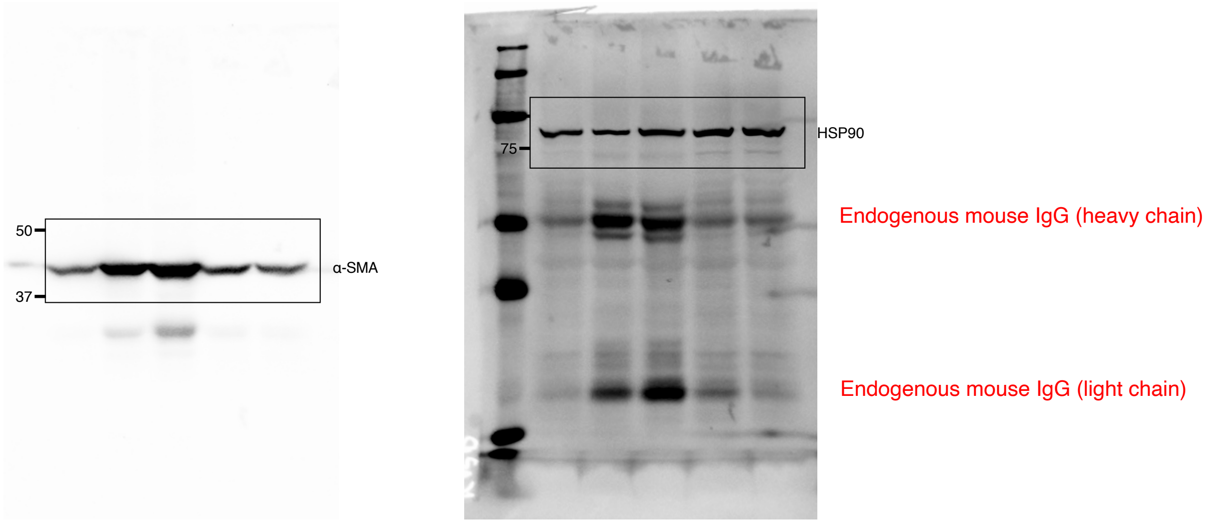


Full-length western blots from Supplemental Figure 5a

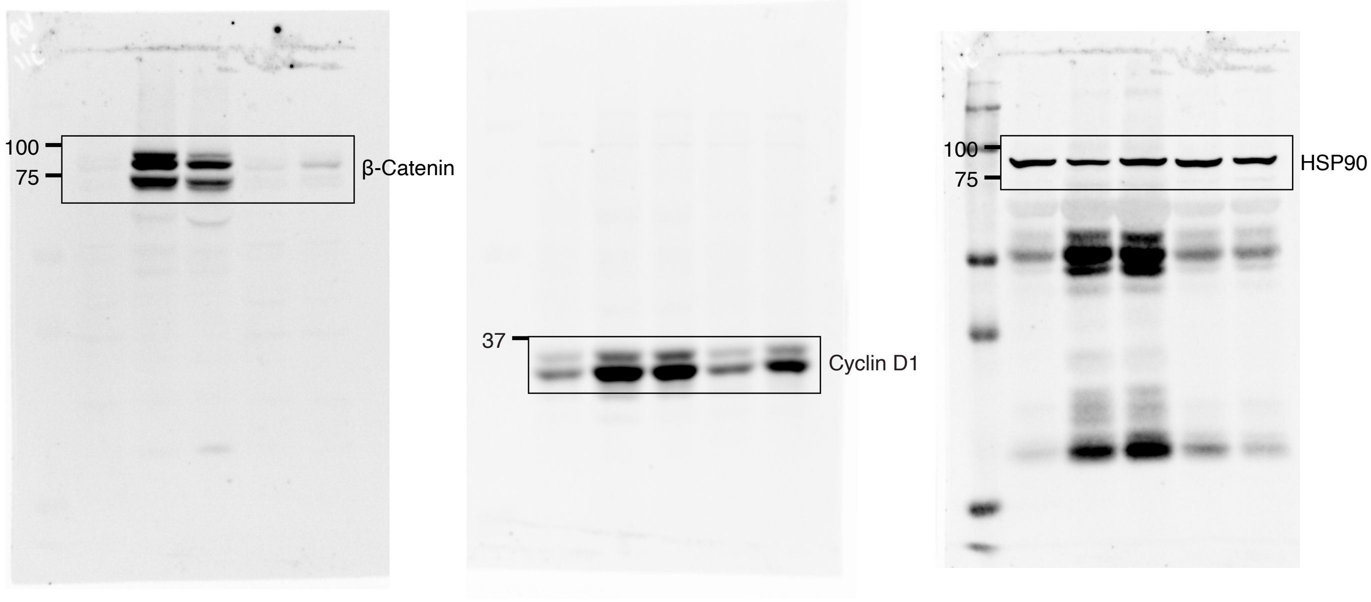

### Full-length western blots from Figure 4b

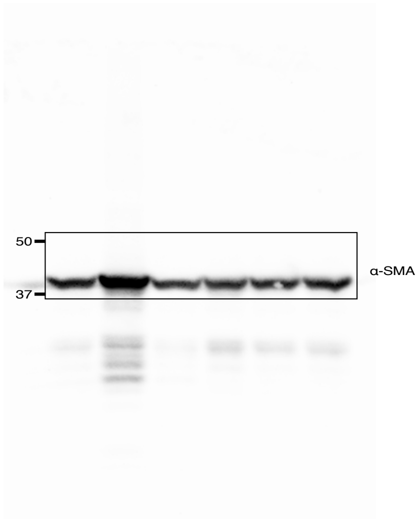


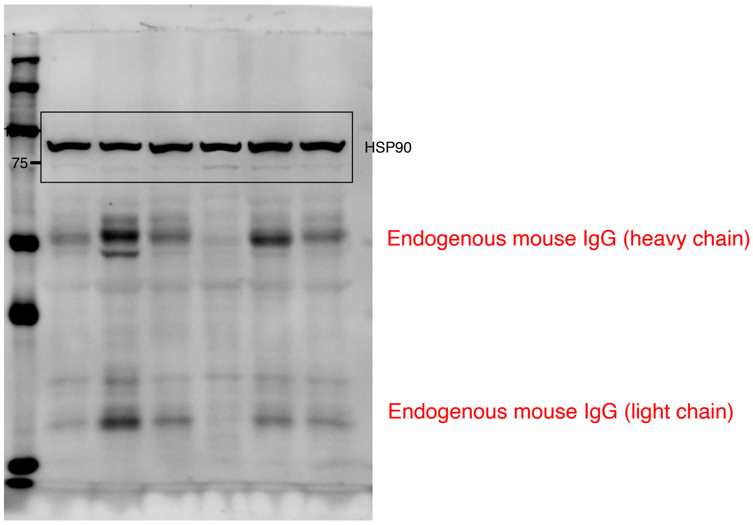


### Full-length western blots from Supplemental Figure 5d

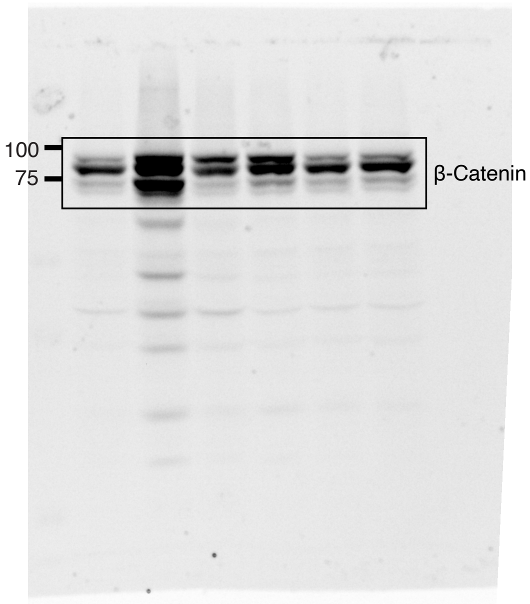


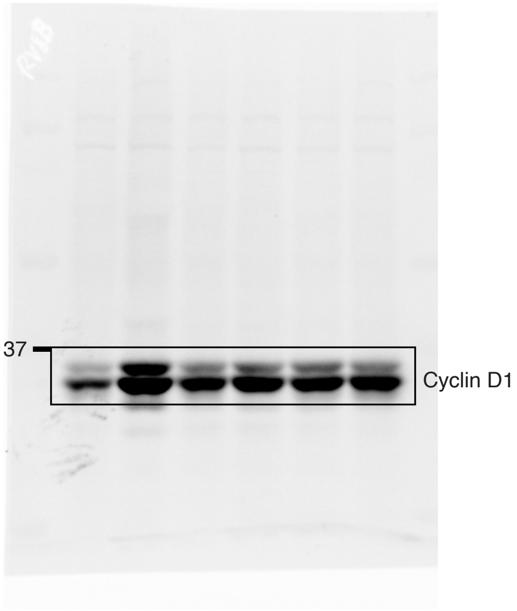


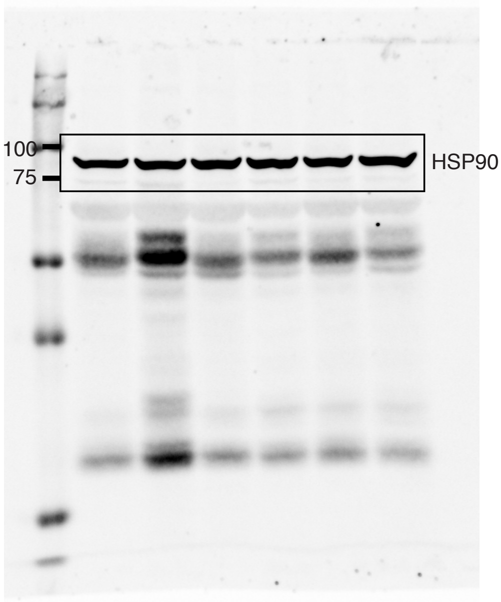
